# Constitutive activation of the PI3K-Akt-mTORC1 pathway sustains the m.3243A>G mtDNA mutation

**DOI:** 10.1101/2020.06.18.159103

**Authors:** Chih-Yao Chung, Kritarth Singh, Vassilios N Kotiadis, Gabriel E Valdebenito, Jee Hwan Ahn, Emilie Topley, Joycelyn Tan, William D Andrews, Benoit Bilanges, Robert D S Pitceathly, Gyorgy Szabadkai, Mariia Yuneva, Michael R Duchen

## Abstract

Mutations of the mitochondrial genome (mtDNA) cause a range of profoundly debilitating clinical conditions for which treatment options are very limited. Most mtDNA diseases show heteroplasmy – tissues express both wild-type and mutant mtDNA. While the level of heteroplasmy broadly correlates with disease severity, the relationships between specific mtDNA mutations, heteroplasmy, disease phenotype and severity are poorly understood. We have carried out extensive bioenergetic, metabolomic and RNAseq studies on heteroplasmic patient derived cells carrying the most prevalent disease related mtDNA mutation, m.3243A>G. These studies reveal that the mutation promotes changes in metabolites which is associated with the upregulation of the PI3K-Akt-mTORC1 axis in patient-derived cells and tissues. Remarkably, pharmacological inhibition of PI3K, Akt, or mTORC1 activated mitophagy, reduced mtDNA mutant load and rescued cellular bioenergetic function. The rescue was prevented by inhibition of mitophagy. The PI3K-Akt-mTORC1 axis thus represents a potential therapeutic target that may benefit people suffering from the consequences of the m.3243A>G mutation.

## INTRODUCTION

Mitochondria control cellular bioenergetic homeostasis and serve as a hub for cell metabolism and cell signalling pathways. Human mitochondria contain a circular plasmid-like DNA (mtDNA) which encodes 13 proteins which act as subunits of the electron transport chain (ETC) and 24 RNAs essential for mitochondrial protein synthesis. Mutations of mtDNA affect around 1 in 5000 of the population^1^ and cause a range of diseases for which no effective treatment is available^2, 3^. Over half of the known pathogenic mutations are found within tRNA genes, in which the m.3243A>G mutation, a tRNA^Leu^ point mutation accounts for about 40% of adult patients of primary mitochondrial diseases^1, 4^. Clinically, signs and symptoms in patients with mtDNA mutations are highly heterogeneous^2, 5^. The tissues primarily affected vary depending on the specific mtDNA mutations, and our understanding of the relationships between mtDNA mutations, disease phenotype and severity is very limited^2, 3^. The majority of diseases caused by mutations of mtDNA are heteroplasmic – tissues express both normal and mutation carrying mtDNA. This is a confounding complication, as disease expression may differ radically between patients with the same mutation but with different mutant load. While disease severity broadly correlates positively with the relative burden of mutant mtDNA, we know remarkably little about the determinants of the mutant mtDNA burden^3, 6–8^.

Mitochondrial quality control pathways, including mitophagy, mitochondrial biogenesis pathways and mitochondrial shaping mechanisms, are critically involved in regulating mitochondrial energy homeostasis^9, 10^. Several studies have demonstrated the accumulation of damaged mitochondria and defective mitophagy as a hallmark of mtDNA diseases and age-related neurodegeneration, suggesting that the presence of pathogenic mtDNA alone is not sufficient to drive selection against the mutation by the activation of mitophagy^11–15^. We therefore asked how cell signalling pathways influence these pathways in the disease model, as adaptive - or maladaptive – responses to impaired oxidative phosphorylation (OxPhos) and changes in intermediary metabolism, and whether these pathways play a role as determinants of mutant load and disease severity.

In this study, we have characterised the metabolic phenotype of patient-derived cells bearing the m.3243A>G (tRNA^Leu^) mutation. This is the most common heteroplasmic mtDNA mutation^1, 4^, and, clinically, is expressed variably but may include diabetes, sensorineural deafness, myopathy, encephalopathy, lactic acidosis and stroke-like episodes (MELAS). We have found that the basal expression and activity of the PI3K-Akt-mTORC1 pathway were increased in the mutant cells, and were strongly associated with redox imbalance, oxidative stress, and glucose dependence. Phosphorylation of Akt and ribosomal protein S6 (a downstream target of mTORC1) were increased in muscle biopsies from other patients with the mutation, confirming that this signalling pathway is constitutively activated in patient tissues. Remarkably, inhibition of PI3K, Akt, or mTORC1 all substantially reduced mutant load and rescued mitochondrial bioenergetic function. This process is mediated by the upregulation of mitophagy which is absolutely required for rescue of mitochondrial function and reduction in mutant load. These findings thus reveal that in response to the m.3243A>G mutation, metabolism is rewired, and activity of the PI3K-Akt-mTORC1 axis is increased, presumably as an adaptive response to metabolic changes driven by the mutation. The finding that inhibition of the pathway reduces mutant load and rescues mitochondrial bioenergetic function suggests that activation of this signalling pathway is, in fact, maladaptive, that activation of the pathway sustains disease progression and that pharmacological intervention in this signalling pathway represents a potential therapeutic strategy in patients with these dreadful diseases.

## RESULTS

### The m.3243A>G mutation causes mitochondrial dysfunction and glucose dependence, resulting in redox imbalance and oxidative stress

To explore the metabolic and cell signalling impact of the m.3243A>G mutation, we used six cell lines: fibroblasts derived from two patients carrying the mutation, two controls matched for age and gender, an A549 cybrid cell line carrying the mutation and its Wild-type (WT) counterpart. PCR-RFLP^16^ was used to ensure the presence of the m.3243A>G mutation and ARMS-qPCR^17^ was used to quantify the m.3243A>G mutation load in these cell models. One patient-derived line (henceforth referred to as patient 1) showed a mutant load of 86.2 ± 2.3%; a second line (referred to as patient 2) showed a mutant load of 30.3 ± 3.5%, and the mutant load was 79.0 ± 0.3% in the cybrid cells (Fig. 1A; for more details about the patients please see Methods).

**Figure 1.**
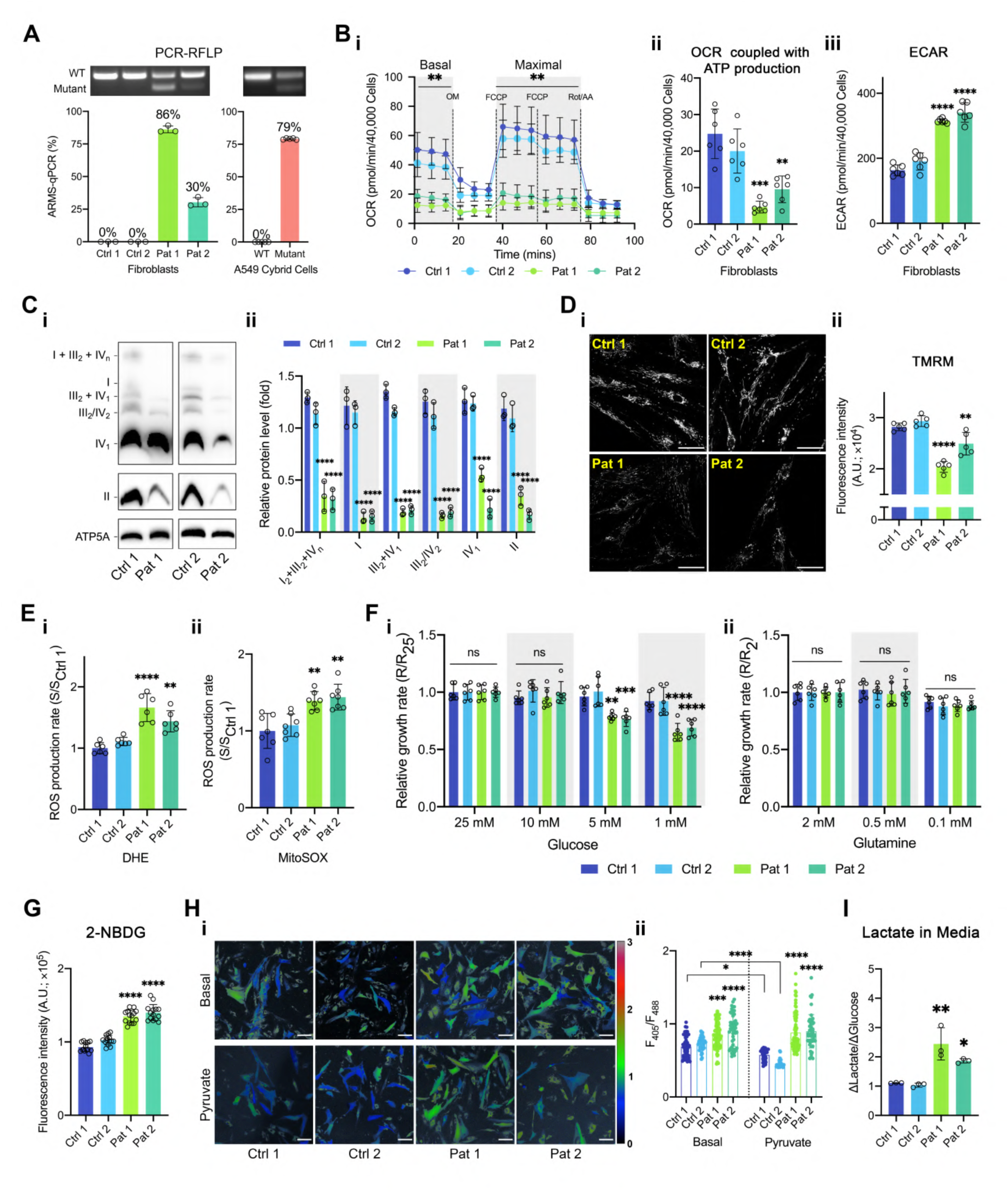
The m.3243A>G mtDNA mutation causes mitochondrial dysfunction and switches cell metabolism to a more glycolytic phenotype, resulting in redox imbalance and oxidative stress. (A) PCR-RFLP and ARMS-qPCR were used to quantify the mutation load for patient-derived fibroblasts (n = 3 independent biological samples) and A549 cybrid cells (n = 5 independent biological samples). (B) Cell respiratory capacity was measured using the Seahorse XFe96 extracellular flux analyser in patient fibroblasts (n = 6 culture wells) showing a major decrease in oxygen consumption under all conditions (i). Oxygen consumption dependent on ATP production (ii) – the response to oligomycin – and extracellular acidification rate (ECAR) are plotted (iii). (C) The expression of respiratory chain proteins and supercomplex assembly in patient fibroblasts were assessed using blue native gels electrophoresis (BNGE, i) and quantified (ii), showing a major decrease in the assembly of all supercomplexes from the patient-derived cells (n = 3 independent experiments) (D) The mitochondrial membrane potential of fibroblasts was measured using TMRM with confocal microscopy (i) and quantified (ii), showing a significant decrease in potential (n = 5 independent experiments). Scale bar = 50 μm. (E) ROS production rates of patient fibroblasts were measured using the reporters, DHE (i) and MitoSOX (ii). Rates of production were significantly increased in patient-derived cells (n = 6 independent biological samples). (F) The growth rates of patient fibroblasts were measured under a range of different nutrient conditions (normalised to the growth rate of each cell line in regular cell media), showing a decrease of growth rate compared to controls at glucose concentrations of 5 and 1 mM (i) but not at low glutamine (ii) concentrations (n = 6 culture wells for all conditions, 3 independent experiments). (G) Glucose uptake in patient fibroblasts was measured using the fluorescent glucose analogue, 2-NBDG, showing a significantly increased rate of glucose uptake in the patient fibroblasts (n = 14 culture wells). (H) NADH:NAD^+^ ratio of fibroblasts was measured using the reporter, SoNar, under basal conditions and after addition of pyruvate (200 μM, 30 min; i) and quantified (ii; n = 50 ± 10 cells). Scale bar = 100 μm. (I) Lactate production (normalized to glucose consumption) was measured in the media of fibroblasts using CuBiAn and showed a significant increase in lactate release in the media of patient-derived cells (n = 4 independent biological samples). All data are represented as mean ± S.D. and data were analysed by one-way ANOVA with Tukey’s multiple comparisons test (* *p* < 0.05, ** *p* < 0.01, *** *p* < 0.001, **** *p* < 0.0001).

To characterise the metabolic phenotype of the mutant cells, respiratory rate was measured using the Seahorse XFe96 extracellular flux analyser. These measurements showed a profound decrease in resting respiratory rate, ATP dependent respiration and in maximal respiratory capacity and an increase in extracellular acidification rate (ECAR) in both patient lines (Fig. 1B). Consistent with these findings, immunoblotting of respiratory chain supercomplexes using blue native gel electrophoresis (BNGE), which identifies non-denatured macromolecular assemblies, revealed disrupted expression of respiratory chain proteins in the mutant cells (Fig. 1C). Notably, there was a very large decrease in assembly of supercomplexes I_2_+ III_2_ +IV_n_, III_2_+IV_1_ and III_2_/IV_2_ and in complex IV_1_. Surprisingly, levels of complex II, which is entirely encoded by nuclear genes, were also consistently and significantly reduced in all mutant cell lines. Mitochondrial membrane potential (Δψ_m_) was reduced in both patient-derived fibroblast lines (Fig. 1D). The decrease in Δψ_m_ was greater in cells from patient 1 than from patient 2, consistent with the greater mutant load in patient 1.

Mitochondrial dysfunction may alter rates of free radical generation by the respiratory chain. The rate of increase of dihydroethidium (DHE) fluorescence (Fig. S1A), reflecting the intracellular rate of production of reactive oxygen species (ROS), was significantly increased both in patient fibroblasts compared to matched controls (Fig. 1Ei). MitoSOX, a mitochondria-targeted form of DHE, revealed a significant increase in the rate of ROS generation in the mitochondria in the mutant cells (Fig. 1Eii), suggesting that the increased rate of ROS generation is likely the consequence of an impaired respiratory chain. Together, the m.3243A>G mutation results in mitochondrial dysfunction and elevated ROS generation.

We wondered whether glucose or glutamine metabolism is altered in the mutant cells, supporting cellular bioenergetic homeostasis and compensating for the mitochondrial dysfunction. Using time-lapse live-cell imaging, we quantified cell growth rates by fitting the curves of cell confluence with an exponential cell growth model (Fig. S1B) under a variety of nutrient conditions. Growth rates were significantly reduced in all mutant fibroblasts compared to their control counterparts (Fig. S1C). Moreover, the cell proliferation rate was significantly reduced in all mutant cells in 1 and 5 mM glucose media (Fig. 1F) but not in low concentrations of glutamine, suggesting that the patient-derived cells are significantly more dependent on glucose metabolism. Consistently, the rate of uptake of the fluorescent glucose analogue, 2-NBDG, was significantly increased in patient fibroblasts compared to controls (Fig. 1G).

The cytosolic NADH:NAD^+^ ratio is a function of glycolytic flux (which consumes NAD^+^, generating NADH), lactate production through lactate dehydrogenase (LDH) which produces lactate and regenerates NAD^+^ from NADH and the activity of malate/aspartate and the glycerol phosphate shuttles that exchange NADH and NAD^+^ between cytosol and mitochondrial matrix. We used the genetically encoded reporter SoNar^18^ to quantify the cytosolic NADH:NAD^+^ ratio (Fig. S1D). The basal NADH:NAD^+^ ratio was significantly increased in the patient fibroblasts compared to controls, consistent with increased regeneration of NADH through LDH activity (Fig 1H). Addition of exogenous pyruvate can either support mitochondrial activity, increasing OxPhos, or stimulate LDH activity to generate lactate, changing the cytosolic NADH:NAD^+^ ratio. In control cells, pyruvate addition decreased the cytosolic NADH:NAD^+^ ratio, but had no significant effect on the high NADH:NAD+ ratio in the mutant cells (Fig. 1H), suggesting that in the latter, LDH flux is already saturated. These data are also consistent with significantly higher levels of lactate produced by the patient fibroblasts compared to controls (Fig. 1I). The pH of the growth media, reflecting cellular lactate production, was also measured using the absorbance of the pH indicator, phenol red. The acidification of media by mutant cells was significantly greater than that of controls (Fig S1E). These data recapitulate the lactic acidosis which is characteristic of patients with the m.3243A>G mutation^19^, and suggest that in patient fibroblasts, increased activity of LDH may compensate for the decreased regeneration of NAD^+^ by mitochondria, supporting increased glycolytic flux. Of note, these metabolic features were recapitulated in the A549 cybrid cells (Fig S1C, F-L), including cell growth rate, OCR (since OCR is affected by cellular mitochondrial content, its value was normalised to their increased mtDNA copy number, Fig. S1F), ECAR, Δψ_m_, ROS generation, glucose uptake, glucose dependence, the cytosolic NADH:NAD^+^ ratio. Together, these data confirm that glucose uptake and glucose catabolism into lactate are upregulated and cytosolic NADH:NAD^+^ ratio is increased in the mutant cells.

### The m.3243A>G mutation remodels glucose metabolism towards increased lipid biosynthesis

To investigate the glucose-dependent metabolic alterations in the mutant cells and identify metabolic signals which might drive changes in cell signalling, glucose uniformly labelled with isotopic carbon ([U-^13^C]-glucose) was used to trace its metabolic fate in the cells by gas chromatography-mass spectrometry (GC-MS). The pattern and abundance of ^13^C enrichment in downstream metabolites were used to determine the relative contribution of glucose into different metabolic pathways. First, partial least squares discriminant analysis (PLS-DA) for the concentration of metabolites (Fig. 2A) showed a clear separation between patient fibroblasts and controls (Fig. S2A). The analysis of metabolic pathways revealed enrichment of glycerolipid metabolism, pentose phosphate pathway (PPP), glycolysis, the tricarboxylic acid (TCA) cycle and aspartate metabolism (Fig. S2B). Tracing the fate of ^13^C-glucose through glycolysis (Fig. 2B), we found that the enrichment of ^13^C in glucose-6-phosphate (G6P) in the patient fibroblasts was at a similar level as in controls, but the total pools of G6P were increased (Fig. 2A) consistent with increased glucose uptake. Although the end products of glycolysis, intracellular pools of pyruvate and lactate in the patient fibroblasts, were maintained at similar levels with controls in terms of ^13^C enrichments and concentrations (Fig. 2A and 2B), the levels and the ^13^C enrichment of glycerol-3-phosphate (αG3P) were increased. The latter suggests the increased activity of glycerol-3-phosphate dehydrogenase in response to increased cytosolic NADH:NAD^+^ ratio, which may drive elevated lipid biosynthesis. Interestingly, while the absolute concentrations of alanine were increased in the mutant cells, its ^13^C enrichment was decreased, suggesting a contribution from other nutrient sources. Similarly, the total concentration of serine was increased in patient fibroblasts, despite decreased synthesis of serine from glucose, suggesting increased serine uptake from extracellular media (Fig. 2A, 2B and S2C).

**Figure 2.**
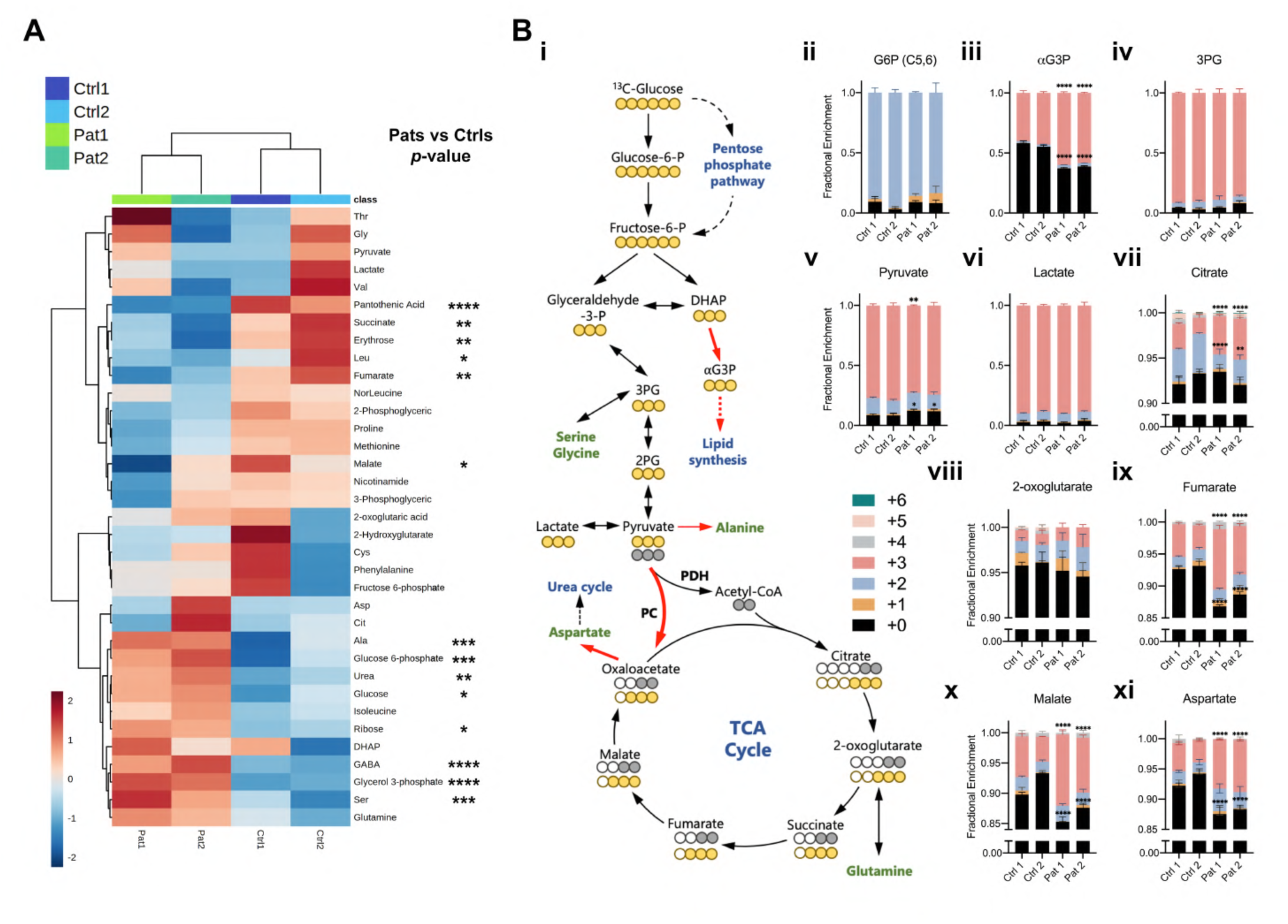
The m.3243A>G mutation rewires glucose metabolism towards increased anabolic biosynthesis. (A) The heatmap of metabolite concentrations obtained by GC-MS was generated by MetaboAnalyst 5.0 (n = 3 technical replicates), revealing significant differences in the concentrations of several metabolites between controls and patient fibroblasts. (B) Schematic of simplified glucose metabolism of the cells (i) Red arrows, upregulated pathways or reactions; three yellow circles of pyruvate and beyond, pyruvate is converted to OAA by pyruvate carboxylase; three or two grey circles of pyruvate and beyond, pyruvate is converted to acetyl-CoA by PDH. Tracing the distribution and abundance of ^13^C enrichment in the intermediates of glycolysis (ii-vi) and the TCA cycle (vii-xi) combined with Fig 2A and S2B revealed an increase in lipid synthesis and PC activity (n = 3 independent biological samples). All data, except Fig. 2A, are represented as mean ± S.D. and were analysed by one/two-way ANOVA with Tukey’s multiple comparisons test for fibroblasts (* *p* < 0.05, ** *p* < 0.01, *** *p* < 0.001, **** *p* < 0.0001).

Consistent with OxPhos defects, levels of the TCA cycle intermediates, malate, fumarate and succinate were decreased in the mutant cells (Fig. 2A). Interestingly, the enrichment of m+3 isotopologues of these TCA cycle intermediates and aspartate were significantly increased in the patient fibroblasts (Fig. 2B). To enter the TCA cycle, glucose-derived pyruvate is converted into acetyl-CoA by pyruvate dehydrogenase (PDH). In this case the catabolism of [U-^13^C]-glucose results in m+2 enrichment of TCA cycle intermediates, including oxaloacetate (OAA), a precursor of aspartate. In contrast, m+3 isotopologues of TCA cycle intermediates are generated if pyruvate is converted into OAA by pyruvate carboxylase (PC). Immunoblotting for PC and phospho-PDH expression revealed elevated PC expression in both patient fibroblasts and cybrid cells and increased levels of phosphorylated PDH (indicating its reduced activity) in patient 2 fibroblasts and cybrid cells (Fig. S2D). As the glucose contribution into the TCA cycle is usually low in conventional cell culture systems, this slight but significant difference in the enrichment of ^13^C is meaningful. The switch from PDH to PC as means of channelling glucose-derived carbon into the TCA cycle in mutant fibroblasts is consistent with the increased NADH:NAD+ ratio, which stimulates the activity of pyruvate dehydrogenase kinase, phosphorylating PDH and inhibiting its activity. Our findings from the stable-isotope-labelling approach and comparative metabolomics thus reveal a profound remodelling of glucose metabolism as a consequence of OxPhos defects arising from the m.3243A>G mutation.

### Chronic activation of the PI3K-Akt-mTORC1 axis in the m.3243A>G mutant cells impairs autophagic flux and mitochondrial degradation via mitophagy

The metabolic phenotype shown above points to reprogramming of nutrient-sensing signalling networks in the m.3243A>G mutant cells. We therefore carried out an RNA sequencing screen to examine global changes in the expression profiles of the patient fibroblasts (Fig. S3A). Using a significance level of false discovery rate (FDR) < 0.05, we identified 3394 transcripts of which 1849 were upregulated and 1545 were downregulated (Fig. S3B). A multi-dimensional principal component analysis also confirmed good accordance between biological triplicates as well as close clustering of transcripts from patient fibroblasts (Fig. S3C). The enrichment analysis identified ‘mitochondrial dysfunction’, ‘TCA cycle’ as the altered metabolic pathways in patient fibroblasts (Fig. S3D). Analysis of individual genes involved in these pathways showed a general downregulation of OxPhos genes (Fig. S3E). Specifically, the increased expression of PC was consistent with the immunoblotting data (Fig. S2D) and correlated with altered TCA cycle activity (Fig. S3F). The enhancement of glycolytic flux in the mutant cells reported by the metabolomic data was supported by the increased expression of hexokinase 1 (HK1). A general decrease of enzymes in serine/glycine metabolism (Fig. S3G) and an altered expression of amino acid transporters (Fig. S3H) also matched our observation of increased serine in the metabolomics measurements. These findings further validated the metabolic footprint observed in patient-derived cells from the metabolomic studies.

Consistent with the above results, the network analysis of these differentially expressed genes identified the strong enrichment of PI3K-Akt and mTOR pathways in patient fibroblasts (Fig. 3A and Fig. S3I; Table S1). We therefore validated these data at the protein expression level. Immunoblotting confirmed increased phosphorylation of Akt (S473) relative to total Akt, as well as phosphorylation of the downstream mTORC1 target S6 ribosomal protein in patient fibroblasts (Fig. 3B) and the cybrid cells (Fig. S3J). While control fibroblasts showed a growth factor-dependent activation of Akt in response to increased serum concentration in growth media, phospho-Akt was elevated in patient cells even when growth factors were low (1% FBS) and increased only slightly in response to an elevated serum concentration. Similarly, the patient fibroblasts exhibited increased mTORC1 activation independently of the serum concentration, revealed by increased phospho-S6 levels. PI3K-Akt-mTORC1 signalling also antagonizes the activity of the cellular energy sensor, AMP-activated protein kinase (AMPK) and suppresses macromolecular catabolism by autophagy. We therefore measured AMPK phosphorylation in response to different serum concentration by immunoblotting, but found no significant difference between patient and control fibroblasts. We further measured phosphorylation of the Akt substrates, p-TSC2 (S939) and p-PRAS40 (T246), and the mTORC1 substrate, p-4EBP1 (S65), of patient fibroblasts grown in 10% or 1% FBS media. In all of these, phosphorylation was increased in cells carrying the mtDNA mutation independent of serum concentration, again consistent with constitutive activation of the pathway in the mutant cells (Fig.S3K). These results suggest that the constitutive induction of PI3K-Akt-mTORC1 signalling remodels cell metabolism, independently of growth factor stimulation.

**Figure 3.**
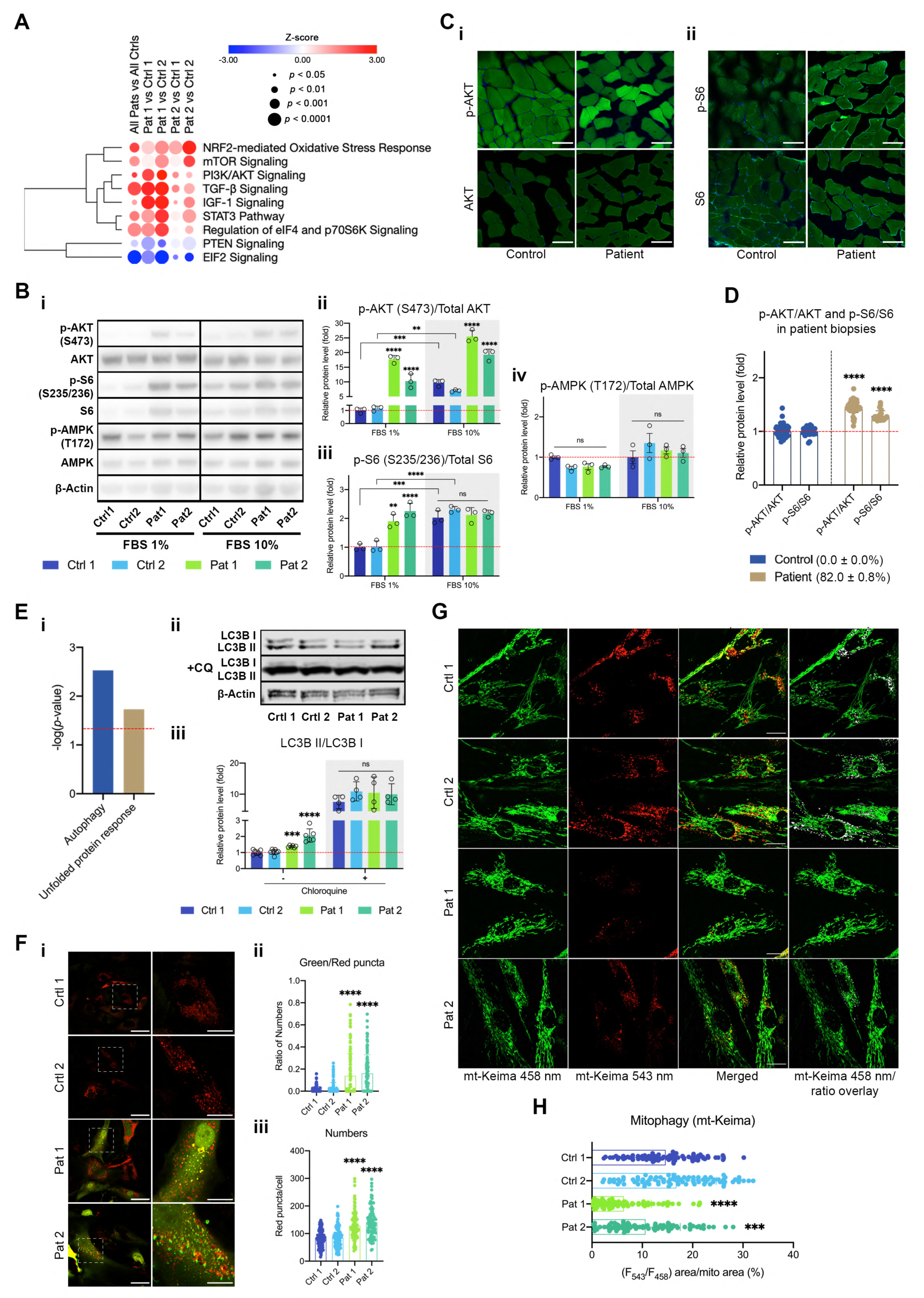
The PI3K-Akt-mTORC1 axis is upregulated in the m.3243A>G mutant cells, perturbing autophagy and mitophagy. (A) Analysis of RNA-seq data from the patient fibroblasts by QIAGEN Ingenuity Pathway Analysis (IPA) showed cell signalling pathways which are enriched in cells carrying the m.3243A>G mutation. These include striking and significant increases in expression of pathways involving PI3K-Akt, mTOR, EIF2, and NRF2. It is notable also that PTEN expression, a negative regulator of PI3K-Akt-mTOR signalling was downregulated (*p* < 0.05). (B) Immunoblotting of p-Akt (S473)/Akt, p-S6 (S235/236)/S6 and p-AMPK (T172)/AMPK in patient fibroblasts grown in the presence of 10% or 1% FBS media (i), showing a major increase in p-Akt/Akt (ii) and p-S6/S6 (iii) in the presence of 1% FBS media in the mutant cells. In contrast, there is no difference in p-AMPK/AMPK (iv) (mean ± S.D. of n = 3 independent experiments). (C-D) Immunofluorescence staining for p-Akt (S473)/Akt (Ci, n > 60 muscle fibres) and p-S6 (S235/236)/s6 (Cii, n > 25 muscle fibres) in patient muscle biopsies confirmed the increased phosphorylation of Akt and S6 in the patients (D). Scale bar = 100 μm. (E) Analysis of RNA-seq data from the patient fibroblasts by IPA showed enriched autophagy and unfolded protein response (i) which is associated with activation of PI3K-Akt-mTOR signaling (see Fig. S4A). Immunoblotting of LC3B with chloroquine (CQ, 50 µM for 5 h; n = 4 independent biological samples) or without CQ (n = 7 independent biological samples) in patient fibroblasts (ii, and quantified in iii) suggests an accumulation of LC3BII in the patient cells without an increase in autophagic flux. (F) Confocal imaging of live cells transfected with the autophagy reporter, mCherry-GFP-LC3 in patient fibroblasts (i, n > 100 cells). The ratio of green/red puncta (ii) and autophagosome numbers (iii) were further quantified, showing an increase of these indices. Scale bar = 50 μm in full-scale and 20 μm in zoomed-in images. (G-H) Live cell imaging of mt-Keima expressing control and patient fibroblasts showing mitophagy events at 543 nm excitation. The overlay of mt-Keima emission at 458 nm and ratio image shows the relative level of mitophagy events in control vs patient fibroblasts. (H) The individual mitophagy events, as represented in ratio image, were quantified and plotted as percentage high (F_543_/F_458_) ratio area/total mitochondria area, showing a significant reduction in ‘mitophagosome’ maturation in patient fibroblasts (n≥ 80-100 cells). Scale bar = 20 μm All data, except Figs. 3A and 3Ei, are represented as mean ± S.D. and were analysed by one-way ANOVA with Tukey’s multiple comparisons test for fibroblasts and by unpaired t test for biopsies (* *p* < 0.05, ** *p* < 0.01, *** *p* < 0.001, **** *p* < 0.0001).

To establish that the PI3K-Akt-mTORC1 axis is also constitutively active in patients with the m.3243A>G mtDNA mutation, we measured p-Akt/Akt and p-S6/S6 by immunofluorescence in a muscle biopsy from a patient with an 82% mutant load of the m.3243A>G mutation causing MELAS, (not one of the donors used to obtain the fibroblasts; matched H&E and COX staining are shown in Fig. S4A). These data confirmed a significant increase in p-Akt and p-S6 in the muscle biopsy, which argues that upregulation of this pathway is a clinical feature of the disease and not a function of the cultured cells (Fig. 3C-D). Re-analysis of a published RNA-seq data set^22^ from muscle biopsies of MELAS patients also confirmed the activation of PI3K-Akt-mTORC1 signalling in patients (Table S4). In summary, activation of the PI3K-Akt-mTORC1 axis is a common feature of m.3243A>G mutation both in vivo and in vitro.

Analysis of the differentially expressed genes in patient fibroblasts also identified autophagy and unfolded protein response as the enriched cellular process in patient fibroblasts (Fig. 3Ei; Table S2-S3). We therefore explored the impact of chronic induction of PI3K-Akt-mTORC1 signalling on autophagic flux in the mutant cells (Fig. S4Bi)^13, 20^. The mRNA expression of genes involved in autophagy was highly altered in patient fibroblasts (Fig S4Bii). We next monitored LC3B turnover in the presence and absence of chloroquine (CQ) or Bafilomycin A1 (Baf A1), both of which prevent lysosomal acidification and inhibit function, using LC3B immunoblotting as a readout. The mutant cells showed an increased conversion to LC3BI to the lipidated LC3BII form (autophagosome) but the difference in LC3BII accumulation in the presence of CQ was not significantly different between control and mutant cells (Fig. 3Eii-iii), indicating a defect in the late stage of autophagosome to autolysosome conversion/fusion in the mutant cells. We also measured the autophagic flux using a mCherry-EGFP-LC3B tandem reporter^21^ by live-cell imaging, in which the EGFP is quenched in an acidic environment, seen as red puncta. Consistent with the above result, the ratio of green/red puncta was low in control compared to patient cells, representing the formation of mature autolysosomes and a high autophagy flux. In contrast, the ratio of green/red puncta and both the number and size of double-positive puncta were increased in mutant cells, indicating impaired autophagosome maturation into autolysosomes (Fig. 3F). These features, LC3B turnover and autophagic flux, were recapitulated in patient fibroblasts treated with Baf A1 (Fig. S4C) and in the A549 cybrid cells treated with CQ or Baf A1 (Fig S4D-F). Altogether, these results confirm that autophagic flux is impaired in the mutant cells and further indicate that an autophagy imbalance may contribute to metabolic and proteostatic stress in the mutant cells as a consequence of the activation of PI3K-Akt-mTORC1.

Given the impaired mitochondrial form and function and autophagy imbalance observed in m.3243A>G mutant cells, we sought to examine the mitochondrial quality control by mitophagy pathway. Similar to mCherry-EGFP-LC3B, we used COX8-EGFP-mCherry (EGFP-mCherry targeted to mitochondrial matrix) as a mitophagy reporter in cybrid cells. The proportion of mitophagic positive cells was quantified by a flow cytometry-based approach in which an arbitrary threshold of red fluorescence was used to indicate the mitophagic polulation. The flow cytometry data revealed an increase in mitophagic population in response to oligomycin and antimycin A (Oligo+AA)-induced mitophagy, which was completely abolished when the control cells were co-treated with Baf A1. However, the mitophagic population in cybrid cell did not alter significantly when treated with either Oligo+AA or Oligo+AA+Baf A1 suggesting an accumulation of autophagosomes-containing mitochondria or its impaired maturation (Fig. S4G). To visualise the delivery of impaired mitochondria to lysosomes, we transfected the fibroblasts with mitochondrial-targeted mKeima (mt-Keima), which undergoes a shift in excitation spectrum within acidic lysosomes indicating the conversion of autophagosome to autolysosomes. A ratio of signal excited at 543nm and 458nm (F_543/_F_458_) provides a quantitative readout of mitophagic activity. The control fibroblasts exhibited punctate structures showing strong mt-Keima signal at 543 nm which indicated that a subpopulation of mitochondria was delivered to lysosomes. Quantification of the ratio image also demonstrated a high ratio (F_543/_F_458_) signal corresponding to increased formation of autolysosomes containing degraded mitochondria. In contrast, the patient fibroblasts exhibited fragmented mitochondrial network throughout the cytoplasm displaying a strong fluorescence at 458 nm but a diminished mt-Keima signal at 543 nm. The subpopulation of mitochondria with high ratio (F_543/_F_458_) signal was significantly decreased which suggested a defect in autolysosome maturation in patient fibroblasts (Fig. 3G-H). Altogether these findings indicate that chronic activation of PI3K-Akt-mTORC1 signalling impairs mitophagy pathway which may lead to accumulation of damaged mitochondria in m.3243A>G mutant cell.

### Pharmacological inhibition of PI3K-Akt-mTORC1 signalling reduces mutant load, rescues mitochondria function and lowers glucose dependence in the m.3243A>G mutant cells

At this point, we asked what the functional consequences of the activation of the PI3K-Akt-mTORC1 axis are and whether it represents an adaptation to impaired OxPhos. We therefore systematically inhibited each component of the axis in turn using well-established pharmacological inhibitors (Fig. 4A)^20^ and explored the consequences for mutant mtDNA burden and the pathophysiological and biochemical features that we have described above.

**Figure 4.**
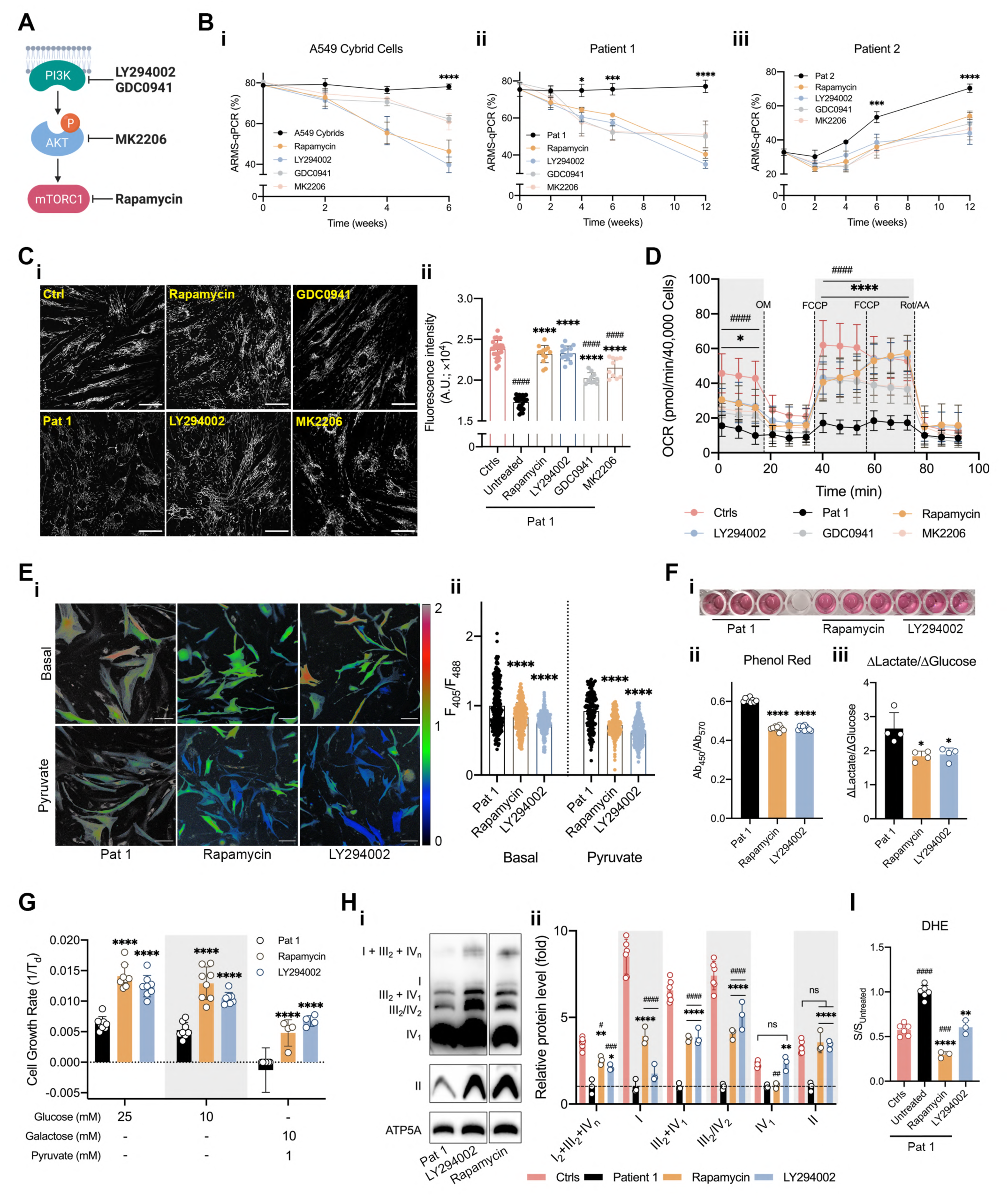
Inhibitors of the PI3K-Akt-mTORC1 axis, LY294002 and Rapamycin, reduce mutation load, rescue mitochondrial function and reduce glucose dependence. (A-B) Schematic depicting pharmacological inhibition of the PI3k-Akt-mTORC1 aixs by LY290042 (LY), GDC0941 (GDC), MK2206 (MK) or Rapamycin (RP) (A). (B) Sustained treatment of cells over 6 or 12 weeks with LY (5 µM) GDC (1 µM), MK (1 µM) or RP (5 µM) caused a progressive decrease in mutant mtDNA load in A549 cybrid cells (i) and fibroblasts of patient 1 (ii) and suppressed the progressive increase of mutant load with time in culture seen in patient 2 fibroblasts (iii) (n = 3 independent experiments). The concentrations of RP and LY here refer to all subsequent treatments. (C) The mitochondrial membrane potential (TMRM 25 nM) of patient 1 fibroblasts treated with LY, GDC, MK or RP for 12 weeks were significantly increased (n = 13 independent biological samples). Scale bar = 50 μm. (D) The respiratory capacity of patient 1 fibroblasts treated with LY, GDC, MK or RP for 12 weeks was measured using the Seahorse XFe96 extracellular flux analyser and showed a major increase in oxygen consumption under all conditions for the drug-treated cells (n = 14 of culture wells). (E) The NADH:NAD^+^ ratio of patient 1 fibroblasts treated with LY or RP for 12 weeks was measured using SoNar under basal condition and pyruvate (200 μM, 30 min; i) and quantified (ii; n > 180 cells). The result showed a significant decrease compared to pre-treatment levels. Scale bar = 25 μm. (F) Acidification of the growth medium revealed by the absorption ratio of phenol red was significantly reduced after chronic drug treatments with RP and LY (i and ii; n = 9 culture wells). Lactate production normalised to glucose consumption (iii, n = 4 culture wells) was also significantly reduced. In each case, patient 1 fibroblasts treated with LY or RP for 12 weeks were treated with fresh media and the media were then collected after 48h incubation. (G) Fibroblasts of patient 1 treated with LY or RP for 12 weeks were also cultured in media with a variety of glucose/galactose concentrations to further assess the glucose dependence, displayed a significant improvement of cell growth in all conditions from the drug-treated cells (n = 4-8 culture wells). (H) BNGE were used to assess the expression of respiratory chain proteins and supercomplex assembly of patient 1 fibroblasts treated with LY or RP (i), showing a major increase in the assembly of almost all supercomplexes from the drug-treated cells (ii, n = 3 independent experiments). (I) Rates of ROS production of patient 1 fibroblasts treated with LY or RP for 12 weeks were significantly reduced to the level that lower than or no difference from controls (n = 3-6 independent experiments). All data represented as mean ± S.D. and were analysed by one-way ANOVA with Tukey’s multiple comparisons test (* *p* < 0.05, ** *p* < 0.01, *** *p* < 0.001, **** *p* < 0.0001, vs Patient/Mutant control; # *p* < 0.05, ## *p* < 0.01, ### *p* < 0.001, #### *p* < 0.0001, vs WT control).

These inhibitors include two inhibitors of class 1 PI3K, LY294002 (LY, 5 μM) and GDC0941 (GDC, 1 μM), an inhibitor of Akt, MK2206 (MK, 1 μM) and an inhibitor of mTORC1, rapamycin (RP, 5 μM). In both the fibroblasts from patient 1 and the cybrid cells, treatment with LY, GDC, MK, or RP, led to a striking and progressive decrease in mutant mtDNA load over 6-12 weeks of sustained treatment in which the mutant load fell from ∼80% to values as low as 40% (Fig. 4Bi-ii). Interestingly, in the absence of the inhibitors, the fibroblasts from patient 2 showed a progressive increase in mutant load with time in culture, rising from ∼30% to around 70% over 3 months (Fig. 4Biii). Treatment with any of the inhibitors significantly suppressed the rate of increase in mutant load decreasing the total change by around 20%. Although the decrease in mutant load took many weeks to develop, changes in protein phosphorylation state were evident within a day of starting treatment (Fig. S5A-C). Accordingly, inhibition of PI3K by LY or GDC or inhibition of Akt by MK2206 reduced Akt and mTOR phosphorylation, while rapamycin inhibited mTOR phosphorylation but had no impact on Akt. None of the treatments had any immediate effect on AMPK phosphorylation. However, AMPK activity gradually increased at later time points after drug treatments, suggesting an induction of catabolic processes such as autophagy and improvement in sensing the bioenergetic status of the m.3243A>G mutant cells^23^.

To investigate whether the decrease in mutant load after drug treatment was reflected by a concomitant change in mitochondrial function, we measured the mitochondrial bioenergetic status and the metabolic phenotype of the patient-derived cells. Mitochondrial membrane potential of the cells treated with RP or LY showed a complete recovery to levels that were not significantly different from control cells (Fig. 4C). Although Δψ_m_ of the cells treated with GDC or MK was higher than the mutant cells, it was still slightly but significantly lower than controls. Similarly, measurements of mitochondrial respiration showed that both basal (50-65% of controls) and maximal respiratory capacity (69-86% of controls) were increased in cells treated with any of the inhibitors (Fig. 4D). Of note, the rescue of mitochondrial bioenergetics by these inhibitors was recapitulated in the A549 cybrid cells (Fig S5D-E).

We then focused on RP and LY, the two most potent inhibitors, for further experiments. The improvement in mitochondrial respiration suggested a metabolic shift towards OxPhos. Measurements of cytosolic redox state using SoNar (Fig. 4E and S5F) showed a significant reduction in basal NADH:NAD^+^ ratio and increased response to pyruvate addition in the long-term treated patient-derived cells, implying a decreased glycolytic flux. Measurements of glucose and lactate concentrations in the media showed a profound decrease in lactate secretion following LY and RP treatments (Fig 4F). The patient fibroblasts were also cultured in media containing glucose or galactose as a primary carbon source to further assess their dependence on glycolysis. LY-and RP-treated cells, but not mutant controls, grew well in the galactose media, indicating a reduced glycolytic dependence after the drug treatments (Fig. 4G). Also, the mutant cells showed an increased supercomplex assembly as well as rescue of individual components of the mitochondrial ETC following all drug treatments (Fig. 4H). These changes were accompanied by a decrease in the rates of ROS production in cells treated with LY and RP (Fig. 4I). These changes in cell metabolism were again recapitulated in the A549 cybrid cells (Fig S5F-I).

Altogether, these data demonstrate that the long-term treatment of the mutant cells with inhibitors of the PI3K-Akt-mTORC1 axis reduces mutant load and reverses the biochemical consequences of the mutation, rescuing mitochondrial function, reducing glucose dependence and lactate secretion. These data strongly argue that the chronic constitutive activation of the PI3K-Akt-mTORC1 axis serves as a maladaptive response in this disease model and helps to sustain the mtDNA mutant load.

### Reduction of the m.3243A>G mutant load by inhibition of the PI3K-Akt-mTORC1 axis is cell-autonomous

The chronic activation of the PI3K-Akt-mTORC1 axis and increased mutant load have deleterious effects on cellular function and amplify the pathophysiological consequences of the mutation. However, this raises questions about the underlying mechanism by which inhibition of the axis reduces mutant load and rescues mitochondrial function. A possible explanation for the response to drug treatments is that inhibition of the signalling pathway may promote the selective growth of cell populations with low mutant load or selective elimination of cells with high mutant loads. We therefore carried out long term measurements of growth rates and cell death (Fig. 5Ai-ii). Cells were pre-treated with either vehicle, LY, or RP for the time periods as specified in Fig. 5B (pretreatment) and then seeded into 96-well plates for measuring cell growth rate and cell death with either vehicle, LY, or RP (treatment). Although treatment with either LY or RP significantly slowed cell proliferation in both fibroblast and cybrid cells at the time point of 0-week pretreatment (Fig. 5B, upper panel), the changes were small. The cell growth rate was stable in cybrid cells treated with LY or RP, suggesting that there was no clonal expansion. In fibroblasts, growth rates of cells were not altered by LY but increased by RP after 4-week pretreatment. However, the growth rate of patient 1 fibroblasts was not further increased by RP after 8-week pretreatment. In contrast, a significant decrease of mutation load was clearly established after 4 weeks of treatment, suggesting that the change of mutation load is independent of the rate of cell growth. On the other hand, cell death in the drug-treated groups were even lower than in the mutant controls, suggesting that the drug treatments were beneficial, and there was no selective killing of mutant cells (Fig. 5B, lower panel). Treating the cybrid cells with GDC or MK showed similar results in terms of cell growth and death (Fig. S6A-B).

**Figure 5.**
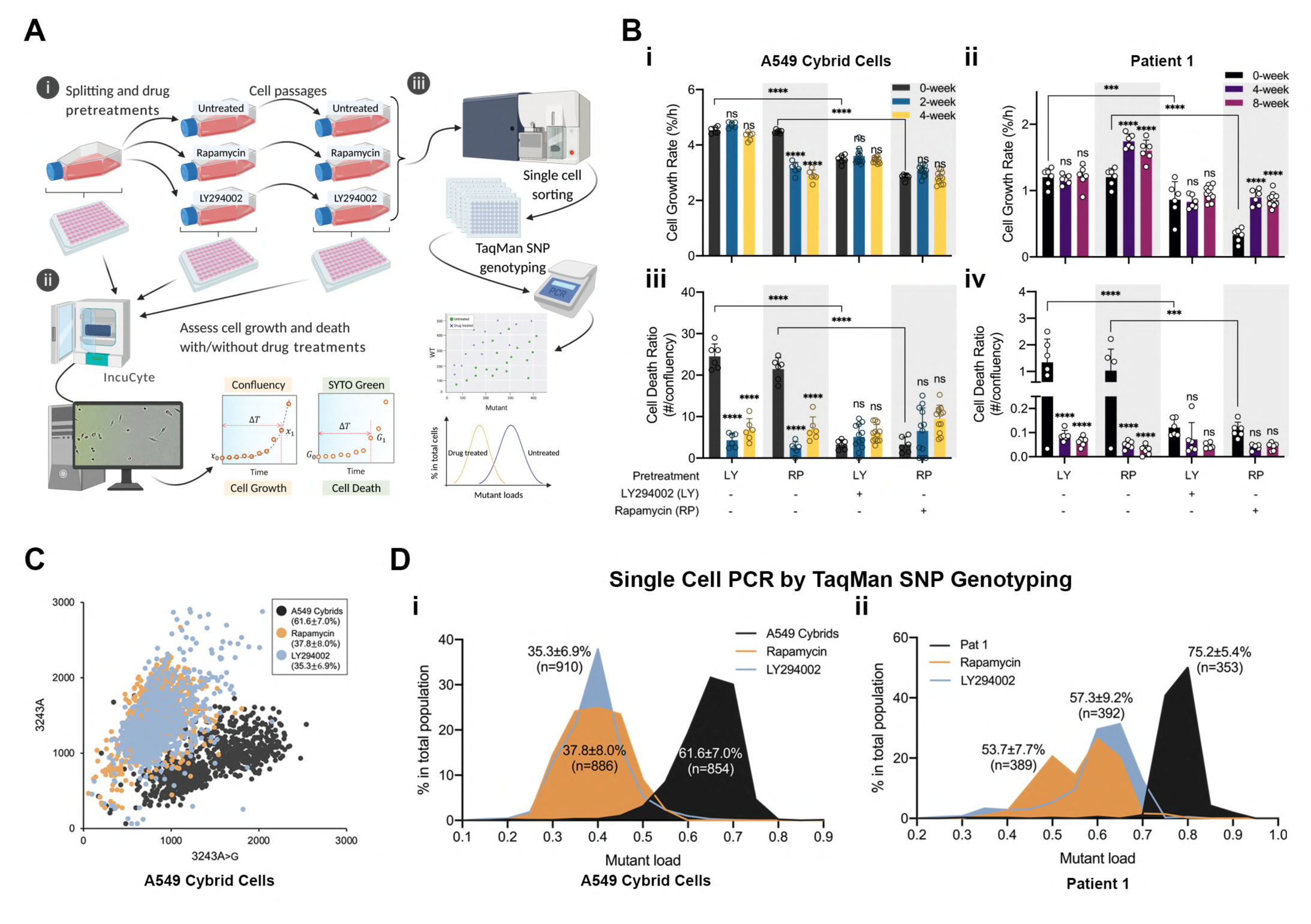
Reduction of the m.3243A>G mutant load by inhibition of PI3K-Akt-mTORC1 requires neither selective cell death nor clonal expansion and is a cell-autonomous event. (A) A flow chart describing the cell culture process (i) used to assess cell growth/death (ii) over the 4-or 8-week drug pretreatments. Cells were pre-treated with either vehicle, LY, or RP for the time periods as specified in Fig. 5B (pretreatment) and then seeded into 96-well plates for measuring cell growth/death with either vehicle, LY, or RP (treatment). Single cell PCR for the m.3243A>G mutant loads was performed at the end of pretreatments using TaqMan SNP genotyping (iii). (B) Cell growth (i and ii) and cell death (iii and iv) over the 4-or 8-weeks drug treatments of A549 cybrid cells (i and iii) and patient 1 fibroblasts (ii and iv) were measured using Incucyte (n = 6-12 culture wells). Although treatment with either LY or RP at 5 μM slowed cell proliferation in both patient 1 fibroblasts and cybrid cells (i and ii), the cell death numbers were also lowered by the drug treatments (iii and iv). (C-D) A scatter plot showing the distribution of WT and mutant mtDNA measured in single A549 cybrid cells treated with LY or RP for 4 weeks showed a clear shift of mutant load distribution. (C). (D) Mutant load distributions of single A549 cybrid cells (i, n > 850 cells) and patient 1 fibroblasts (ii n > 350 cells) treated with LY or RP for 4 and 8 weeks, respectively. All data represented as mean ± S.D. and were analysed by one/two-way ANOVA with Tukey’s multiple comparisons test (* *p* < 0.05, ** *p* < 0.01, *** *p* < 0.001, **** *p* < 0.0001).

To determine definitively whether the reduction of mutant load in response to the drug treatments is cell-autonomous, we used a PCR-based technique, TaqMan SNP genotyping, at single cell resolution to measure the distributions of the m.3243A>G heteroplasmy in a cell population (Fig. 5Aiii). The TaqMan technique was first validated by measuring the mutation loads of a population of cells and then to establish the range of single cell mutant load across the population (Fig. S6C). The distribution of mutant load in the cybrid cells ranged from 50-80%, showing a Gaussian distribution (Fig. S6D). Although we found that this technique is less accurate than ARMS-qPCR and tends to overestimate the mutant load when it is lower than 50% and underestimate the mutant load when it is higher than 50%, the average single cell mutant load was consistent with that of the whole cell population, suggesting a good precision of the technique. Measurement of the relative m.3243A>G heteroplasmy in patient fibroblasts and cybrid cells following treatment with LY and RP for 6 and 12 weeks respectively showed a frequency distribution which clearly segregated the treated group as a distinct population from the untreated group (Figs. 5C-D). The distribution of the mutant load in treated groups shifted to significantly lower levels compared to the untreated group.

To determine whether the reduction in the m.3243A>G mutant load is phenocopied at the protein expression level, we immunostained the mtDNA-encoded cytochrome c oxidase I (MT-COI) in patient 1 fibroblasts. The presence of 7 UUA-encoded leucine residues within MT-COI renders the m.3243A>G tRNA^Leu(UUR)^ cells prone to codon-specific translational defect^24^. Immunofluorescence staining revealed a significant decrease in MT-COI expression in patient 1 fibroblasts compared to controls (Fig. S6E). Following drug treatments, these cells showed a partial but significant recovery of MT-COI expression. The single cell analysis of MT-COI expression segregated the drug-treated and untreated groups in a distribution pattern similar to the m.3243A>G mtDNA distribution shown in Fig. 5D. A diminished expression of MT-COI also correlated with the reduced assembly and activity of supercomplexes shown in Fig. 4H. Together, these results strongly suggest that rather than the enrichment of a specific cell population, the reduction in m.3243A>G mtDNA mutant load caused by inhibition of PI3K-Akt-mTORC1 signalling operates across the whole population and is cell-autonomous.

### Restoration of autophagic/mitophagic flux by PI3K-Akt-mTORC1 inhibition is necessary to reduce the m.3243A>G mutant load

The data from the single cell analysis strongly argue that the change in mutant load is a cell-autonomous phenomenon. We then asked what mechanisms select against the mutant mtDNA. A possible explanation could be the elimination of the mutant mtDNA by reducing the total mtDNA copy number while amplifying the dominant mtDNA, resembling ‘the bottleneck’ effect – the selection mechanism of mtDNA in the germline or embryo^4, 6^. We therefore quantified the total mtDNA copy number of the drug-treated cells at a series of time points, but found no significant changes between the mutant control and drug-treated cells (Fig. S7A) and the total mtDNA copy number remained stable across the duration of drug treatments.

It also seemed plausible that the removal of dysfunctional mitochondria via mitophagy may reduce the mutant load in cells with the mtDNA mutation^25, 26^. Indeed, long-term treatment with LY or RP all significantly increased the conversion and accumulation of LC3BII (autophagosome) in the presence of CQ compared to untreated cells (Fig. 6A). Notably, the autophagic flux, evident by a high LC3BII/LC3BI ratio in the presence of CQ, increased progressively with time closely mirroring the rate of decrease in mutant load (see Fig. 4B). Measurement of autophagic flux using the mCherry-GFP-LC3 probe showed a significant increase in the formation of autolysosomes (red puncta) as early as 24 h of drug treatment and increased upon long-term treatment in patient fibroblasts (Fig. 6B). Moreover, the ratio of green/red puncta numbers decreased significantly, suggesting the restoration of autophagic flux in the m.3243A>G mutant cells after drug treatment. Moreover, similarly results were also found in the cybrid cells and the mutant cells treated with GDC or MK (Fig. S7B-E). Of note, analysis of the cybrid cells transfected with mCherry-GFP-LC3 by flow cytometry also showed an increase in cells with low GFP intensity after RP or LY treatment (Fig S7D).

**Figure 6.**
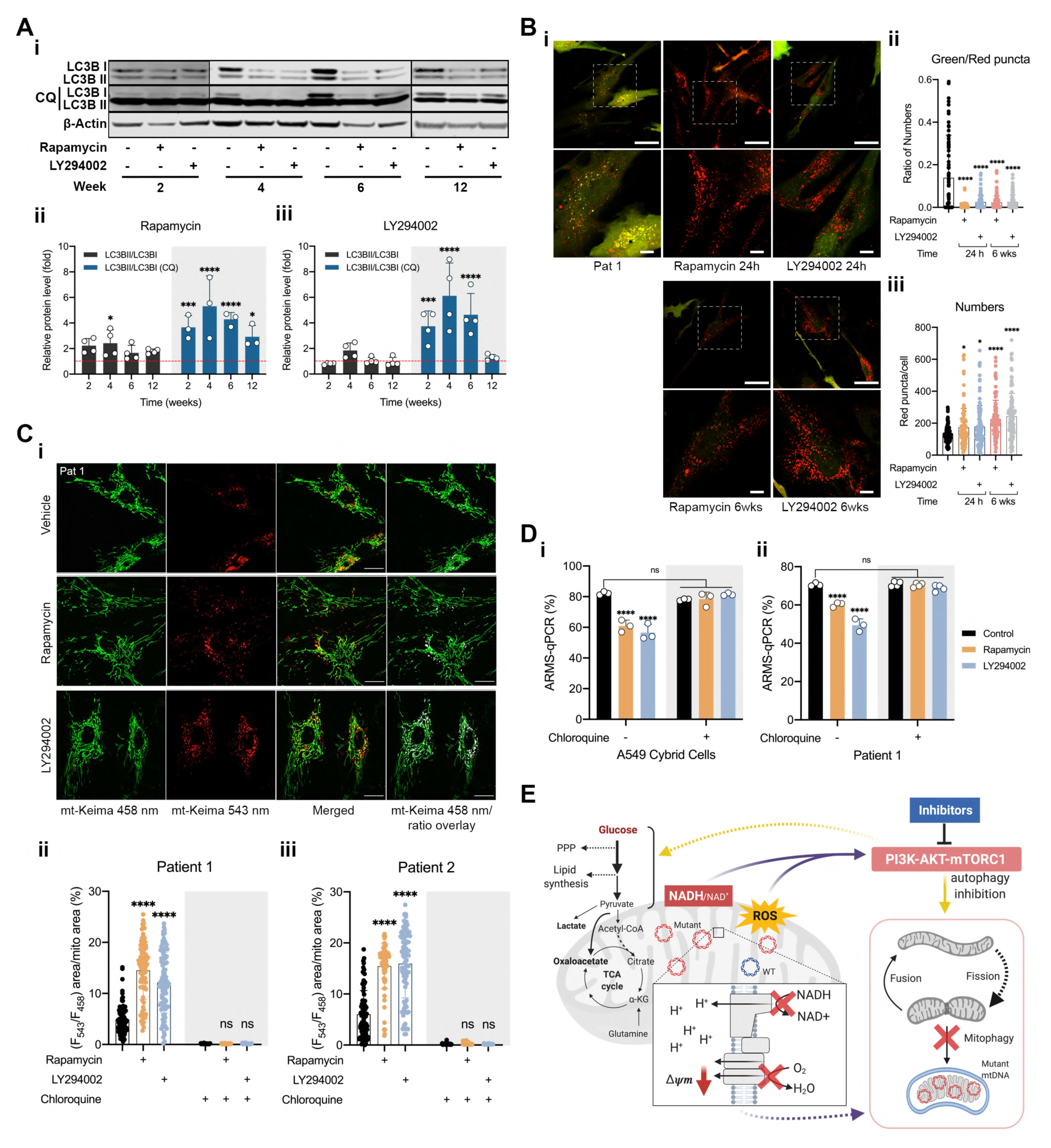
Autophagy/mitophagy is necessary to reduce the m.3243A>G mutant load following inhibition of the PI3K-Akt-mTORC1 axis. (A) In fibroblasts of patient 1, immunoblotting (i) of LC3B with or without CQ (50 μM for 5 h on the day of experiments) over 12 weeks drug treatments (ii and iii) showed a major increase in autophagic flux in drug-treated cells (n = 3-4 of independent experiments). (B) Confocal imaging of the autophagy reporter, mCherry-GFP-LC3 (n > 100 cells) in patient 1 fibroblasts treated with LY or RP at the indicated time points (i, 6 weeks, and ii, 24 h). The ratio of green/red puncta (ii) and autophagosome numbers (iii) were further quantified, showing a major decrease in the ratio of green/red puncta in response to the treatments. Scale bar = 50 μm in full-scale and 10 μm in zoomed-in images. (C) Imaging of the mitophagy reporter (i), mt-Keima, was used to specifically quantify mitophagy in patient fibroblasts treated with LY or RP. Quantification of the mitochondrial area fraction engulfed in autolysosomes in both patient 1 (ii) and 2 (iii) were quantified and showed a dramatic increase in mitophagy in both patient cell lines when treated with RP ad LY (n = 82-95 cells), which was completely prevented by treatment with CQ (n = 32-57 cells). (D) A549 cybrid cells (i) and patient 1 fibroblasts (ii) were cultured with the inhibitors RP and LY as in all prior experiments but in combination with CQ (10 μM), showing that CQ completely prevented the decrease of mutant load in response to inhibition of the PI3K-Akt-mTORC1 pathway (n = 3 independent experiments). (E) A scheme describing how the m.3243A>G mutation alters cell metabolism, including mitochondrial dysfunction, altered glucose metabolism, redox imbalance and oxidative stress, which leads to the constitutive activation of the PI3K-Akt-mTORC1 pathway and a defect in autophagy/mitophagy. Pharmacological inhibition of this pathway reduces mutant load, rescues mitochondrial function and reduces glucose dependence. These findings suggest that the activation of the PI3K-Akt-mTORC1 pathway drives a positive feedback cycle, which maintains/increases mutant mtDNA, augments the metabolic rewiring and thus worsens cell signalling perturbation. This positive feedback loop may shape the phenotype of the disease and determine disease progression. All data are represented as mean ± S.D. and were analysed by one/two-way ANOVA with Tukey’s multiple comparisons test (* *p* < 0.05, ** *p* < 0.01, *** *p* < 0.001, **** *p* < 0.0001).

To evaluate how the increased efficacy of autophagy affects specific status of mitophagy, the cybrid cells were transfected with COX8-mCherry-EGFP, as a reporter of mitophagy. The long-term drug treatment with RP and LY dramatically increased the proportion of cells with low GFP intensity, suggesting the restoration of mitophagy flux (Fig. S7F). Similarly, after 24 h of drug treatments in patient fibroblasts transfected with mt-Keima (Fig. 6C), the (F_543_/F_458_) ratio increased significantly with an increase in red puncta (degraded mitochondria). Altogether, these findings confirm that inhibition of PI3K-Akt-mTORC1 signalling restores autophagy and promotes mitophagic flux in the m.3243A>G mutant cells.

The progressive decrease in mutant load and increase in autophagic flux with time following PI3K-Akt-mTORC1 inhibition in the m.3243A>G mutant cells strengthened the idea that upregulated mitophagic flux drives selection against the mutant mtDNA and decreases the mutant load. To explore this further, we quantified mitophagy in patient fibroblasts expressing mt-Keima, cotreated with CQ in combination with RP or LY. The cotreatment with CQ reversed the effect of LY and RP on mitophagy, decreasing the high ratio (F_543_/F_458_) signal to levels comparable to untreated patient fibroblasts (Fig. 6C and S7G). Remarkably, CQ treatment over 6 weeks in the patient fibroblasts and cybrid cells completely prevented the decrease of mutant load in response to PI3K-Akt-mTORC1 inhibition (Fig. 6D), demonstrating that increased mitophagy is necessary to reduce the burden of mutant mtDNA. This also argues strongly that the reduction in mutant load must be a cell-autonomous effect and cannot be attributed to the selective growth of subpopulations of the cells. To confirm the mechanisms are specific to the m.3243A>G mutation, we also examined the effects of LY and RP in cells carrying another heteroplasmic mtDNA point mutation – the m.8993T>G – generated by the Minczuk lab^27^. In these cells, there is no difference in the phosphorylation of Akt and S6 and autophagic flux assessed by LC3B (Fig. S7Hi). Furthermore, inhibition of the PI3K-Akt-mTORC1 axis had no impact on mutant load (Fig. S7Hii), suggesting that hyperactivation of the PI3K-Akt-mTORC1 axis is relatively disease specific. Together, effective autophagy/mitophagy triggered by inhibition of the PI3K-Akt-mTORC1 axis is necessary to eliminate the mutant m.3243A>G mtDNA.

## DISCUSSION

The clinical presentation of diseases caused by pathogenic mtDNA mutations is highly heterogeneous^4–6^. The association between specific mtDNA mutations, heteroplasmic mutant load, disease manifestation and severity are poorly understood. In the present study, we have systematically characterised the metabolic and cell signalling phenotype of cells bearing the m.3243A>G mutation, the most prevalent mtDNA mutation, which is responsible for mitochondrial encephalomyopathy, lactic acidosis and stroke like episodes (MELAS) or Maternally inherited Diabetes and Deafness (MIDD) and related disorders. We found that the changes in metabolism were associated with the constitutive hyperactivation of the PI3K-Akt-mTORC1 pathway specifically to the m.3243A>G mtDNA mutation. Most notably, pharmacological inhibition of PI3K, Akt, or mTORC1 in the patient-derived cells reduced mutant load and rescued mitochondrial bioenergetic function. These inhibitors promoted mitophagy, which was absolutely required to mediate the effects of these treatments (Fig. 6E). In contrast to the m.3243A>G mtDNA variant, previous studies have reported downregulation of Akt activity in cells carrying the m.8993T>G mutation^28^, in which inhibition of the PI3K-Akt-mTORC1 pathway had no impact on mutant load or mitochondrial function (Fig. S7H). These findings suggest that the changes in signalling pathways are disease specific and may represent a link between genotype and phenotype.

Metabolic features of the patient fibroblasts and cybrid cells are consistent with most of the previous studies of the m.3243A>G mutation, including mitochondrial dysfunction, upregulated glycolysis, increased ROS production, and redox imbalance^5, 15, 19, 23, 24, 29^. However, metabolic profiling of cells bearing the mutation is limited in the literature. Metabolomics data showed that reductive carboxylation of glutamine is a major mechanism to support cell survival and maintain redox balance in the m.8993T>G mutation cell models^27, 30^. In contrast, using a similar approach, we found that glucose anabolism (i.e. upregulated upper glycolysis, PPP, and lipid synthesis) is increased in the m.3243A>G mutant cells and these cells are dependent on glucose for cell survival and proliferation. Also, increased TCA cycle entry of pyruvate through PC, which, combined with the prediction of enriched malate-aspartate shuttle pathway (Fig. S2B) may serve as a mechanism to maintain cellular NADH:NAD^+^ balance to support glycolytic flux and promote antioxidant capacity^27, 31^. This difference in metabolism between the m.8993T>G and m.3243A>G mutations argues that the mtDNA mutations cannot be seen as one disease^32, 33^. Thus, the rewired metabolism specific to the m.3243A>G mutation drew our attention to the corresponding changes in cell signalling.

The alterations of metabolism in the m.3243A>G mutant cells strongly imply a perturbation in cell signalling, especially implicating the PI3K-Akt-mTORC1 pathway, which has been widely studied in cancer signalling and metabolism^34^. For example, oxidative stress and redox imbalance may activate PI3K-Akt, the increased glycolytic flux and PPP implies an activated Akt-NRF2 signalling, the increased PC expression and p-PDH suggests an accumulation of acetyl-CoA associated with Akt-mTORC1 activity^34^. Metabolic pathway analysis using RNA-seq data recapitulated these metabolic changes in the patient fibroblasts. Moreover, network analysis using the RNA-seq data further validated the link between the metabolic changes and altered cell signalling in cells carrying the m.3243A>G mutation.

Indeed, the network analysis also identified EIF2 (eukaryotic initiation factor 2) and PTEN (phosphatase and tensin homolog) signalling as the top downregulated pathways (Fig. 3A). PTEN is a negative regulator of the PI3K-Akt-mTORC1 axis. Although the mechanisms of crosstalk between PI3K-Akt-mTORC1 and EIF2 pathways are still unclear, these two pathways seem to communicate and determine cell fate under stress^35, 36^. Of note, IPA upstream analysis showed an upregulation of transcription factors in the EIF2 pathway – ATF4 and DDIT3 (CHOP) (Table S3-4). ATF4 acts as a prototypical downstream target coupling mitochondrial proteotoxic stress to the activation of ISR (integrated stress response)^37^ and derives remodelling of one-carbon metabolism (serine/glycine and folate metabolism)^38^. Although we observed an increase of serine concentration in fibroblasts bearing the m.3243A>G mutation, incorporation of ^13^C-glucose into serine was reduced. Also, expression of genes related to serine/glycine metabolism was generally reduced (see Fig. S3G). These data imply that in fibroblasts carrying the m.3243A>G mutation, unlike ‘Deletor’ mice^38^, serine metabolism is downregulated. On the other hand, given that the UPR^mt^ (mitochondrial unfolded protein response) maintaining mutant mtDNA in a *C. elegans* model is mitopahgy independent^39^, exploring the role of ISR/UPR^mt^ in the maintenance of the m.3243A>G mutation and disease progression by suppressing eIF2α phosphorylation will be interesting. Furthermore, in a published RNA-seq data set of muscle biopsies from patients with MELAS ^22^, FGF21 (∼48 fold) and GDF15 (∼9 fold) were upregulated, matching previous findings reported from muscle tissue of ‘Deletor’ mice^38^. In contrast, in the present study, while we did not find changes in these two genes in the patient fibroblasts, we did find upregulation of FGF16 (∼33 fold), suggesting that changes in FGF pathways in response to mitochondrial dysfunction might be tissue specific.

Previous studies have suggested that different levels of heteroplasmy of the m.3243A>G mutation in cybrid cells result in different gene expression patterns and even in discrete changes in metabolism^40, 41^. Another study showed that JNK (c-Jun N-terminal kinases) was activated by ROS in cybrid m.3243A>G cells, further reducing RXRA (retinoid X receptor alpha) expression^42^. In contrast, our data did not show inhibition of the RXRA pathway and suggest that chronic activation of the PI3K-Akt-mTORC1 pathway is a general response in cells with the m.3243A>G mutation despite very different levels of mutant load. Remarkably, we found that prolonged treatment with inhibitors of each component of the PI3K-Akt-mTORC1 axis reliably and reproducibly reduced mutant burden and rescued mitochondrial function in the mutant cells. Although inhibition of PI3K-Akt-mTORC1 axis has been shown to be beneficial in several mitochondrial and neurological disease models, the underlying mechanisms remain elusive ^13, 20, 43–45^. As a result, the therapeutic efficacy of the inhibition may be distinct and limited in different models. A study of a mouse model of Leigh syndrome showed that rapamycin, while delaying the progression of the disease, failed to improve OxPhos^46^. Similarly, another study showed that mTORC1 inhibition failed to improve either mitochondrial bioenergetics or survival in a mouse model of mitochondrial encephalomyopathy (Coq9^R239X^)^47^. Our data from cells with the m.8993T>G mutation suggest that changes in signalling pathways may differ between mtDNA diseases, perhaps pointing towards the mechanisms that define differences in disease phenotype between different mitochondrial diseases, and emphasising that therapeutic options should be considered separately for each disease related to mtDNA mutations. However, since both the PI3K-Akt-mTORC1 axis and the accumulation of mtDNA mutations have been associated with neurodegeneration, ageing and cancers, our study may implicate in a broader biomedical context^4,6,9,20,34^.

Overall, the m.3243A>G mutation causes a profound cell metabolic and signalling remodelling, although the causality between the observed alterations in metabolism and cell signalling of the m.3243A>G mutant cells is not yet conclusive. Our working model is that activation of the PI3K-Akt-mTORC1 pathway drives a metabolic rewiring of the cell, redirecting glycolytic metabolism and promoting lactate production. Activating the pathway suppresses mitophagy, allowing the mutation to propagate and thus maintaining the mutant load. Inhibition of the PI3K-Akt-mTORC1 axis promotes mitophagy, reduces mutant load and improves bioenergetic competence in a cell-autonomous way. Thus, we propose that nutrient-sensing pathways that may have evolved as adaptive responses to altered cellular metabolism prove to be maladaptive, driving a positive feedback cycle that may shape the phenotype of the disease and determine disease progression. In conclusion, our results suggest that activation of the PI3K-Akt-mTORC1 axis by changes in intermediary metabolism controls the heteroplasmic burden of the m.3243A>G mutation and may define disease progression and severity in a cell-autonomous manner. These data strongly suggest that cell signalling pathways activated by altered metabolism represent potential therapeutic targets that may benefit people suffering from diseases caused by mtDNA mutations.

---

**Figure S1.**
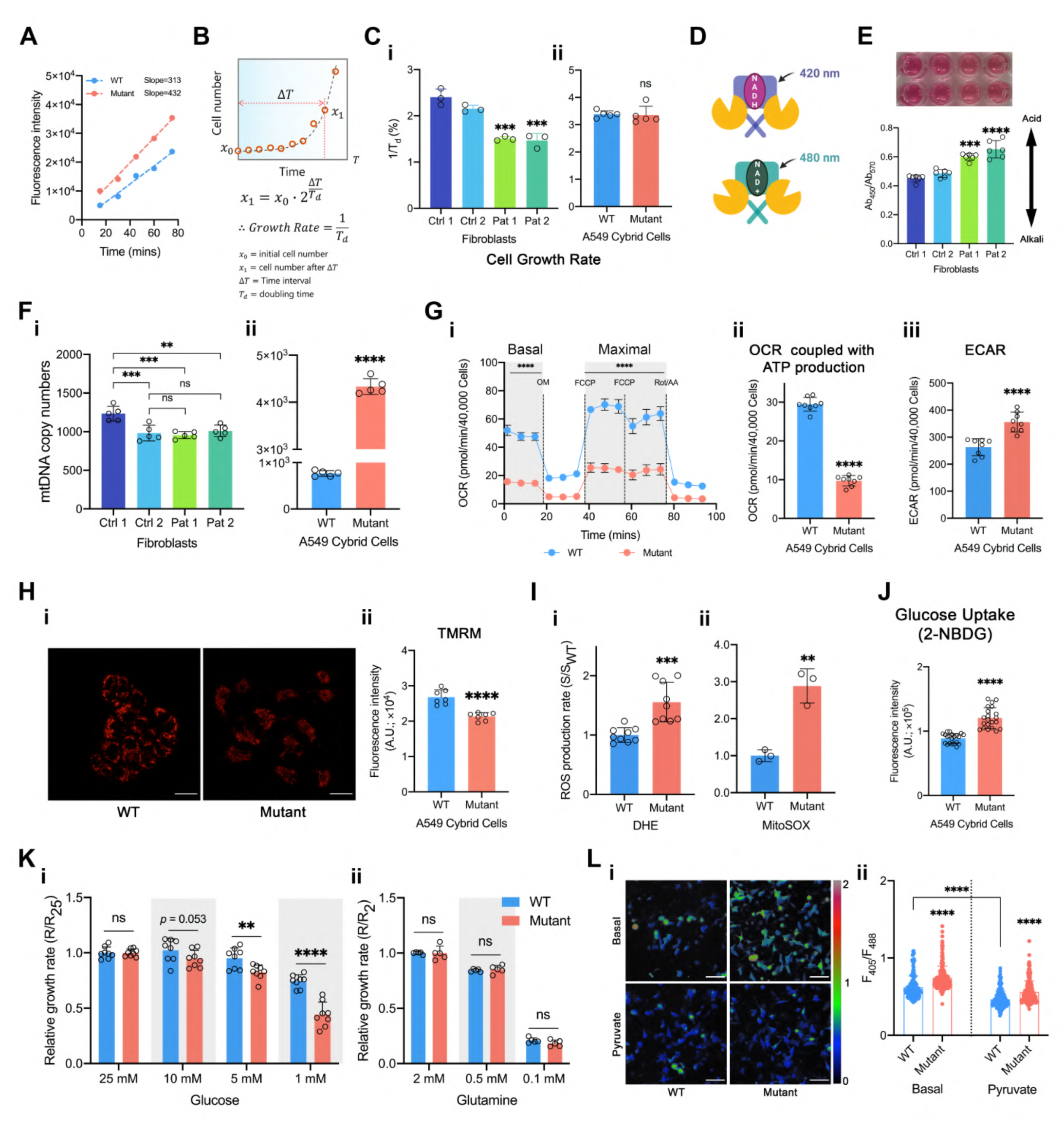
The metabolic phenotype of A549 cybrid cells recapitulates the metabolic features of patient fibroblasts. (A) Rates of ROS production in A549 cybrid cells and the parental cells were measured using dihydroethidium (DHE), as the rate of increase in red fluorescence intensity over 80 min incubation with DHE. (B-C) A scheme describing how growth rates were obtained by fitting growth curves with an exponential cell growth model (B). (C) Based on the model, growth rate of fibroblasts (i, n = 3 culture wells) and A549 cybrid cells cultured with regular medium were obtained (ii, n = 5 culture wells, *p* = 0.84). (D) Schematic depicting the ratiometric probe for NADH:NAD^+^, SoNar. (E) pH in the media from patient fibroblasts cultured for 2 days was measured based on the ratiometric property of the pH indicator, phenol red, in media and showing a lower pH in the media of patient fibroblasts than that of controls (n = 6 independent biological samples). (F) Mitochondrial DNA copy number of fibroblasts (i) and A549 cybrid cells (ii, n = 5 independent bio logical samples). (G) Cell respiratory capacity was also measured using the Seahorse XFe96 extracellular flux analyser in A549 cybrid cells (n = 8 culture wells, further normalised to mtDNA copy number) showing a major decrease in oxygen consumption under all conditions. Oxygen consumption dependent on ATP production (ii) and ECAR are plotted (iii). (H) The mitochondrial membrane potential of A549 cybrid cells measured using TMRM with confocal imaging (i) and quantified (ii, n = 7 independent biological samples). Sale bar = 20 μm. (I) ROS production rates of A549 cybrid cells reported by DHE (i, n = 9 independent biological samples) and MitoSOX (ii, n = 3 independent biological samples). (J) Glucose uptake in A549 cybrid cells was measured by 2-NBDG, showing a significantly increased rate of glucose uptake in the cybrid cells compared to A549 controls (n = 21 culture wells). (K) Cell growth rates of A549 cybrid cells were measured under a range of different nutrient conditions (normalised to the growth rate of each cell line in regular cell media), showing a decreased rate of growth of A549 cybrid cells compared to controls at glucose concentrations of 5 and 1 mM (i; n = 10 culture wells) but not at low glutamine concentrations (ii, n = 6 culture wells). (L) NADH:NAD^+^ ratio of A549 cybrid cells under basal condition and after addition of pyruvate (200 μM, 30 min; i) was measured by the probe and quantified (ii; n > 180 cells). Scale bar = 100 μm. All data are represented as mean ± S.D. and were analysed by one-way ANOVA with Tukey’s multiple comparisons test for fibroblasts and by unpaired t test for cybrid cells (* *p* < 0.05, ** *p* < 0.01, *** *p* < 0.001, **** *p* < 0.0001).

**Figure S2.**
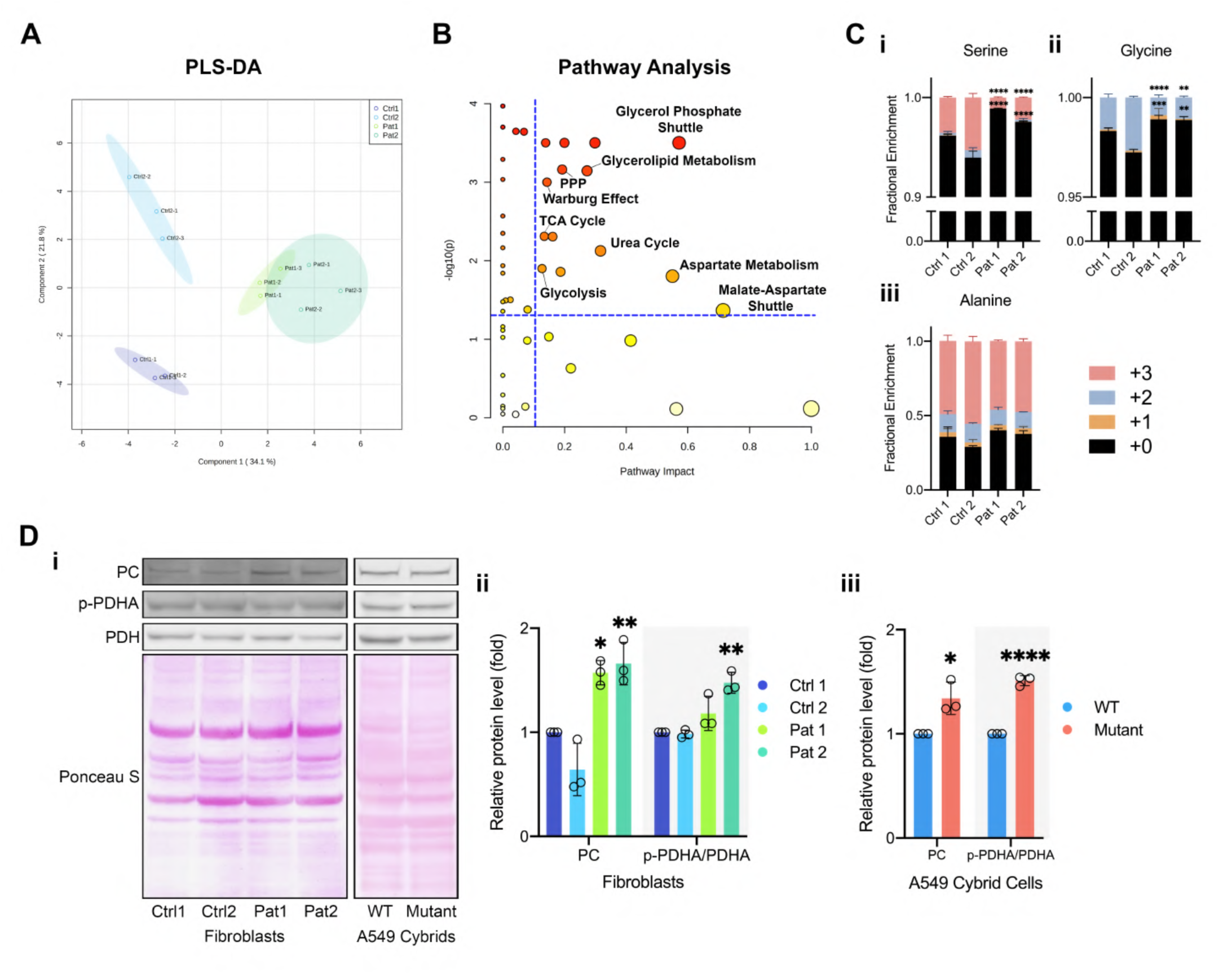
The metabolomic of fibroblasts carrying the m.3243A>G mutation are distinct from that of their matched controls. (A) PLS-DA for the dataset of Fig. 2A showed that the metabolic profiles of patient fibroblasts were distinct from that of controls. (B) The concentration of metabolites obtained by GC-MS was used to determine the enriched metabolic pathways in patient fibroblasts by MetaboAnalyst 5.0., suggesting that phospholipid biosynthesis, PPP, glycolysis, TCA cycle, etc. were enriched (n = 6 technical replicates). (C) With an increase of serine and alanine concentration (Fig 2A) in the patient cells, the incorporation of ^13^C into serine (i) and glycine (ii) were reduced, while alanine (iii) was not altered (mean ± S.D, of n = 3 independent biological samples). (D) Immunoblotting (i) of the expression of pyruvate carboxylase and phosphorylation state of PDH showed an increase of pyruvate carboxylase expression and increased phosphorylation of PDH (S293) in patient fibroblasts (ii) and A549 cybrid cells (iii) (n = 3 independent experiments for all cell lines). All data, except Figs. S2A and S2B, are represented as mean ± S.D. and were analysed by one-way ANOVA with Tukey’s multiple comparisons test for fibroblasts and by unpaired t test for cybrid cells (* *p* < 0.05, ** *p* < 0.01, *** *p* < 0.001, **** *p* < 0.0001)

**Figure S3.**
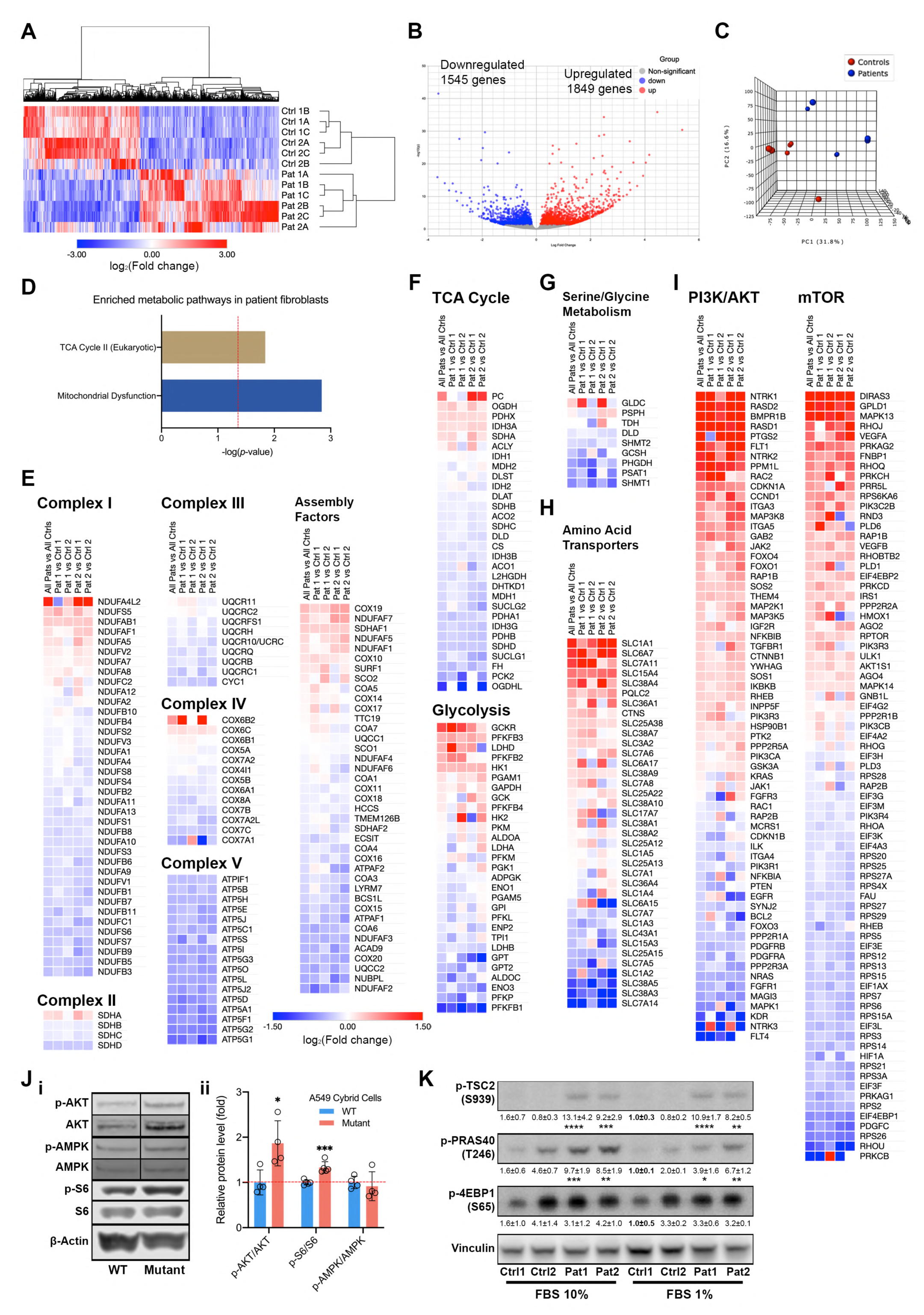
Analysis for RNA-seq of patient fibroblasts with other supporting results and the enriched metabolic pathways concordant with the metabolic phenotype of patient fibroblasts. (A-C) RNA-seq showed by a heatmap (A), a volcano plot (B) and 3D-PCA (C). Gene expression in the mutant cells was significantly distinct from both of the controls: 3394 genes were differentially expressed (FDR < 0.05, with 1849 up-regulated and 1545 down-regulated) (D) Analysis of RNA-seq data from the patient fibroblasts by QIAGEN Ingenuity Pathway Analysis (IPA) showed enriched metabolic pathways (*p* < 0.05) that matchies the findings shown in Figs. 1 and 2. (E) Analysis of the RNA seq data showing a general decrease in mRNA expression of OxPhos-related genes in patient fibroblasts. (F) The mRNA expression of TCA cycle and glycolysis genes in patient fibroblasts confirmed our findings in Fig 2, such as increased pyruvate carboxylase and HK1 expression. (G-H) Analysis of the RNA-seq data showing the mRNA expression associated with amino acid metabolism, including serine/glycine metabolism (G) and amino acid transporters (H). (I) Detailed analysis of the mRNA expression of multiple genes involved in the PI3K-Akt and mTOR pathways in patient fibroblasts, showing consistent differences in a wide array of genes involved in these pathways. (J) Immunoblotting of p-Akt (S473)/Akt, p-S6 (S235/236)/S6 and p-AMPK (T172)/AMPK in A549 cybrid cells (i). (ii) Quantitation shows increased phosphorylation of Akt and S6 but not of AMPK, consistent with the results of fibroblasts (mean ± S.D. of n = 4-5 independent experiments, unpaired t test). (K) Immunoblotting of Akt substrates, p-TSC2 (S939) and p-PRAS40 (T246), and of another mTORC1 substrate, p-4EBP1 (S65), in patient fibroblasts grown in the presence of 10% or 1% FBS media shows increased phosphorylation (n = 3 independent experiments, one-way ANOVA with Tukey’s multiple comparisons test). All data were analysed as stated above (* *p* < 0.05, *** *p* < 0.001)

**Figure S4.**
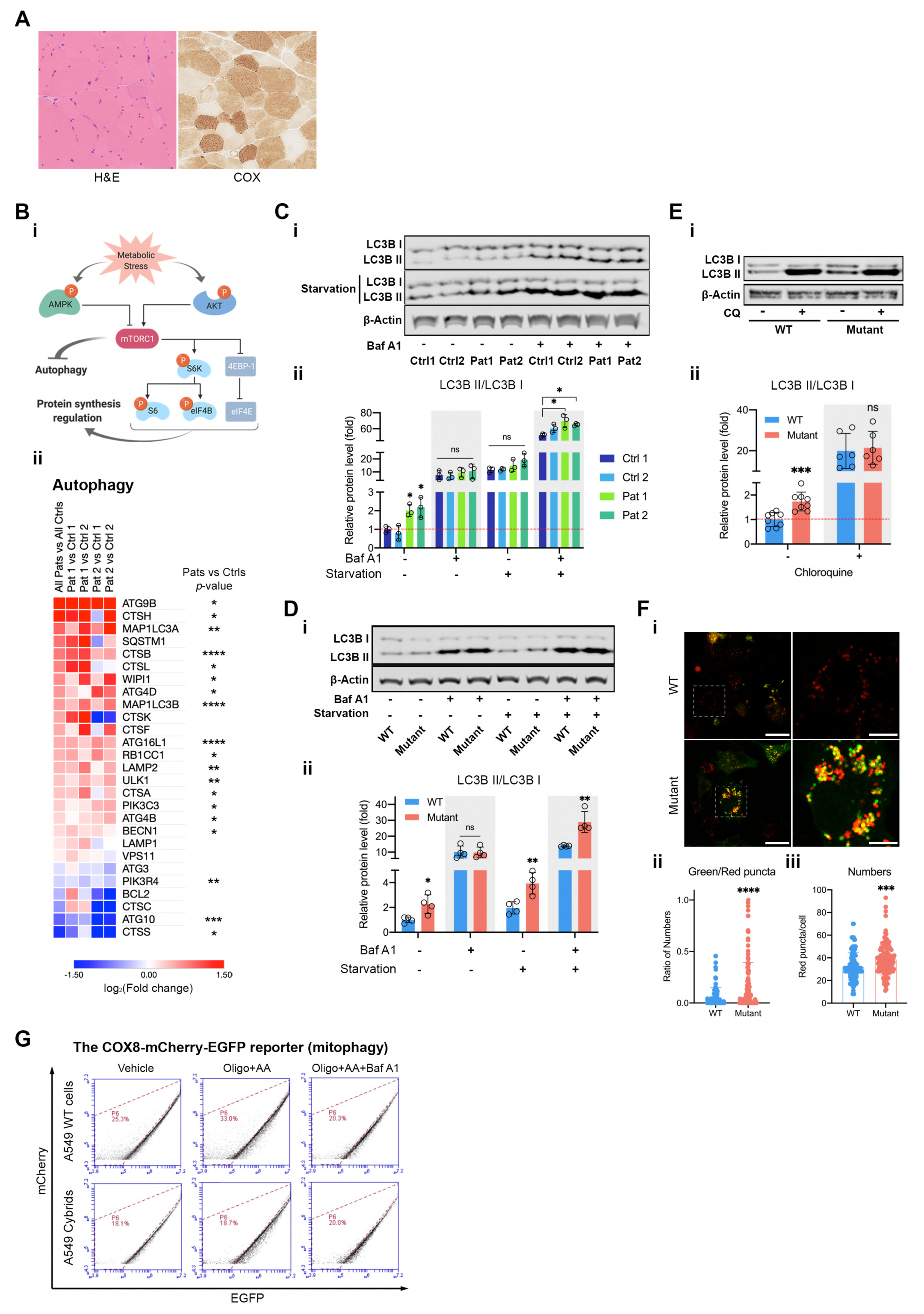
Other supporting results from patient fibroblasts and A549 cybrid cells for assessing autophagy and mitophagy. (A) H&E and COX staining of patient biopsy. (B) A simple scheme showing the regulation of autophagy and protein translation by the PI3K-Akt-mTORC1 pathway (i). Detailed analysis of the mRNA expression of multiple genes involved in the autophagy in patient fibroblasts, showing consistent differences in a wide array of genes involved (ii). (C-D) Immunoblotting of LC3B ± Bafilomycin A1 (Baf A1, 100 nM) in patient fibroblasts (Ci) and in A549 cybrid cells (Di) under basal or starvation conditions (quantified in ii, n = 3-4 independent biological samples) also shows an accumulation of LC3BII in the patient cells without an overall fold increase in autophagic flux that is the conversion of LC3BI to LC3BII. (E) Immunoblotting of LC3B with CQ (50 µM for 5 h; n = 6 independent biological samples) or without CQ (n = 8 independent biological samples) in A549 cybrid cells (i) suggests an accumulation of LC3BII without an increase in autophagic flux (ii) as seen in the patient fibroblasts. (F) Imaging of mCherry-GFP-LC3 (n > 100 cells) in A549 cybrid cells (i). The ratio of green/red puncta (ii) and autophagosome numbers (iii) were further quantified, showing an increase of green/red puncta and total numbers in the mutant cells. Scale bar = 30 μm in full-scale and 10 μm in zoomed-in images. (G) Quantitative analysis of basal autophagy and mitophagy in A549 cybrid cells transfected with COX8-EGFP-mCherry under varied conditions was assessed by flow cytometry using red-to-green fluorescence (n > 3 independent biological samples). All data, except Fig. S4A, are represented as mean ± S.D. and were analysed by one-way ANOVA with Tukey’s multiple comparisons test for fibroblasts and by unpaired t test for cybrid cells (* *p*<0.05, ** *p*<0.01, *** *p*<0.001, **** *p*<0.0001).

**Figure S5.**
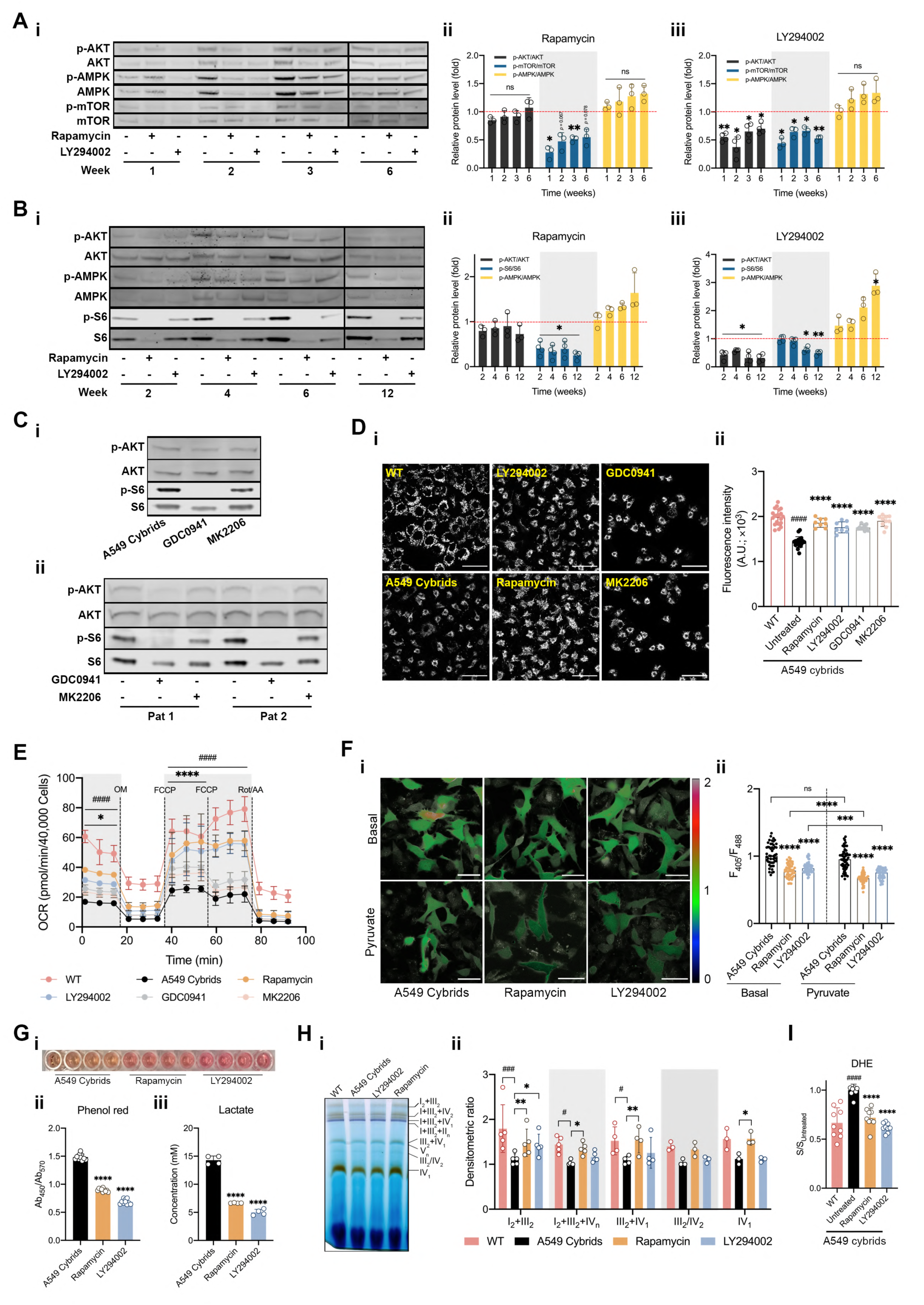
Inhibitors of the PI3K-Akt-mTORC1 axis, LY294002 and Rapamycin, also reduced mutant load and rescued mitochondrial function in A549 cybrid cells. (A) Immunoblotting of the phosphoproteins (p-Akt/Akt, p-mTOR/ mTOR and p-AMPK/AMPK; i) in A549 cybrid cells treated with RP (ii) or LY (iii) over 6 weeks demonstrate the effective inhibition of Akt or mTORC1 (n = 3 independent experiments). (B) Immunoblotting of the phosphoproteins (p-Akt/Akt, p-S6/S6 and p-AMPK/AMPK; i) in patient 1 fibroblasts treated with RP (ii) or LY (iii) over 12 weeks demonstrated the effective inhibition of Akt or mTORC1 in the drug-treated cells (n = 3-4 independent experiments). (C) Immunoblotting of the phosphoproteins (p-Akt/Akt and p-S6/S6) in A549 cybrid cells (i) and in patient fibroblasts (ii) treated with GDC or MK over 6 weeks demonstrated the effective inhibition of Akt or mTORC1 and upregulated autophagic flux in the drug-treated cells (n = 3 independent experiments). (D) The mitochondrial membrane potential of A549 cybrid cells (i, TMRM 25 nM) significantly increased after exposure to LY, GDC, MK or RP for 6 weeks to the level that only slightly lower than which of WT (ii, n = 8 independent biological samples). Scale bar = 25 μm. (E) Cell respiratory capacity of A549 cybrid cells treated with LY or RP for 6 weeks was measured using the Seahorse XFe96 extracellular flux analyser and showed a major increase in basal and maximum OCR after treatment (n = 18 culture wells). (F) Cytosolic NADH:NAD^+^ ratio of A549 cybrid cells transfected with the genetically encoded probe SoNar and treated with LY or RP for 6 weeks was measured (i) and quantified (ii) under basal condition and following exposure to pyruvate (200 μm, 30 min; n > 50 cells). (G) The absorption ratio of phenol red was used to measure the pH of the growth media (i and ii, n=10 culture wells) and the kit of the CuBiAn instrument was used to measure lactate production (iii, n = 4 culture wells). Both acidification of the medium and lactate secretion (48h incubation) of A549 cybrid cells treated with LY or RP for 6 weeks were reduced. (H) BNGE were used to measure supercomplex assembly and In Gel activity of A549 cybrid cells after 6 weeks of treatment with LY or RP (i) and quantified (ii, n = 3 independent experiments). (I) Rates of ROS production of A549 cybrid cells treated with LY or RP for 6 weeks were reduced to the level that no longer significantly different from WT cells (n = 9 culture wells). All data represented as mean ± S.D. and were analysed by one/two-way ANOVA with Tukey’s multiple comparisons test (* *p* < 0.05, ** *p* < 0.01, *** *p* < 0.001, **** *p* < 0.0001, vs Patient/Mutant control; # *p* < 0.05, ## *p* < 0.01, ### *p* < 0.001, #### *p* < 0.0001, vs WT control).

**Figure S6.**
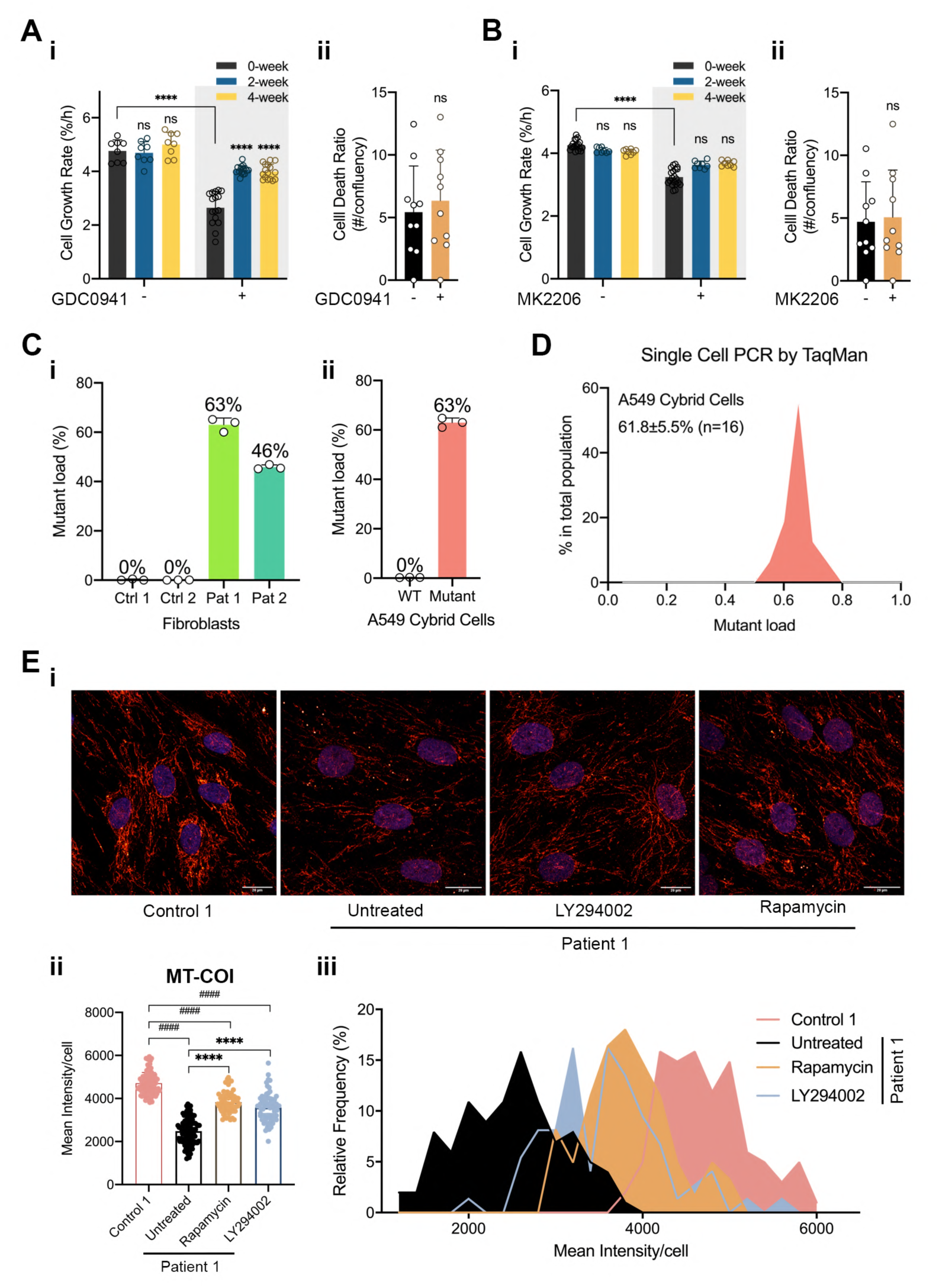
TaqMan SNP genotyping can be used at single cell resolution to measure the distribution of mutant mtDNA burden in a cell population. (A-B) Cell growth and cell death of A549 cybrid cells were measured before and after treatment with GDC (A, n = 8-16 culture wells) or MK (B, n = 8-20 culture wells) for 4 weeks. The drugs slight inhibited cell growth but had no specific effect on the cell death of mutant cells. (C-D) TaqMan SNP genotyping was validated by measuring the mutation load in a whole population of cells (C, n = 3 independent biological samples) and then to establish the range of mutant load at the level of single cells in the A549 cybrid cells (D). (E) Immunofluorescence staining for MT-COI (i and quantified in ii) showed that the expression of MT-COI in patient 1 fibroblasts was significantly lower than that of control 1 (n > 100 cells), while LY or RP treatments restored its expression (n > 60 of cells). The histograms (iii) display the heterogeneous distribution of intensities measured at the single cell level in control 1 and patient 1 fibroblasts. All data are represented as mean ± S.D. and were analysed by one/two-way ANOVA with Tukey’s multiple comparisons test (* *p* < 0.05, ** *p* < 0.01, *** *p* < 0.001, **** *p* < 0.0001).

**Figure S7.**
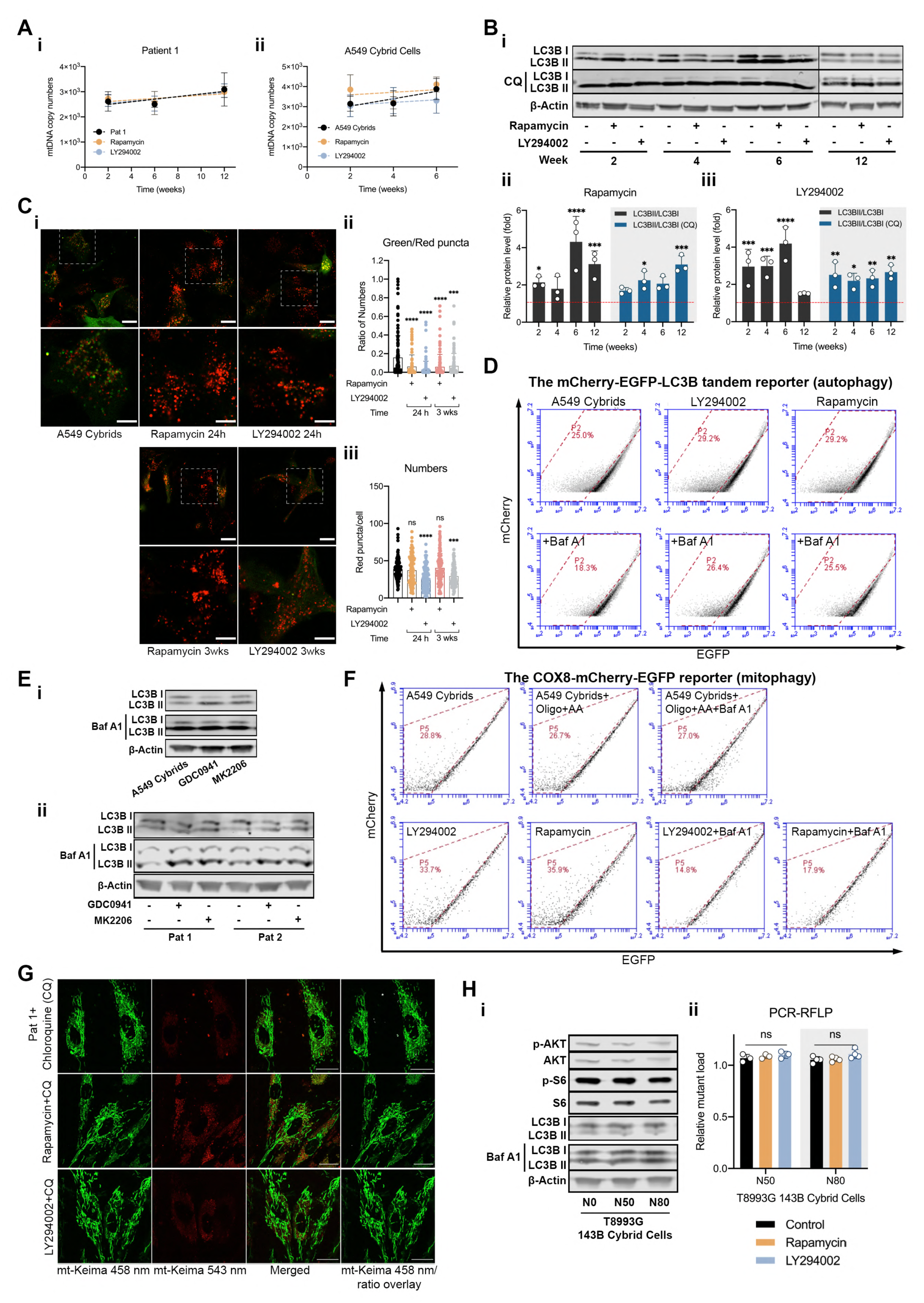
Mitochondrial biogenesis and changes in mtDNA copy number do not explain the reduction of mutant mtDNA burden, while autophagy/mitophagy is also upregulated by inhibition of the PI3K-Akt-mTORC1 pathway in A549 cybrid cells. (A) Quantitative PCR assessing mtDNA copy numbers of patient 1 fibroblasts (i) and A549 cybrid cells (ii) over the 6- or 12-weeks of treatments (n = 3 independent experiments). (B) In A549 cybrid cells, immunoblotting of LC3B with or without CQ (50 μm for 5 h on the day of experiments) throughout drug treatments (i) showed a major increase in autophagic flux in drug-treated cells (ii and iii, n = 3 independent experiments). (C) Imaging of mCherry-GFP-LC3 (n > 100 cells) in A549 Cybrid cells treated with LY or RP at the indicated time points (i). The ratio of green/red puncta (ii) and autophagosome numbers (iii) were further quantified, showing a major decrease in the ratio of green/red puncta and puncta size. Scale bar = 20 μm in full-scale and 10 μm in zoomed-in images. (D) Analysis of A549 cybrid cells transfected with mCherry-GFP-LC3 under varied conditions (n = 3 independent biological samples for each condition) using flow cytometry showed an increase of cells with low GFP intensity when treated with RP and LY, while Baf A1 treatment reversed the effect. (E) Immunoblotting of LC3B (± Baf A1) in A549 cybrid cells (i) and in patient fibroblasts (ii) treated with GDC or MK over 6 weeks demonstrated the upregulated autophagic flux in the drug-treated cells (n = 3 independent experiments). (F) Analysis of A549 cybrid cells transfected with COX8-EGFP-mCherry under varied conditions (n = 3 independent biological samples for each condition) using flow cytometry showed an increase in proportion of mitophagic positive cells in response to LY, RP and Oligo+AA-induced mitophagy while co-treatment with Baf A1 reversed the effect. (G) Imaging of mt-Keima showing that the increased of mitophagy in patient fibroblasts treated with RP or LY was completely prevented by CQ much higher than patient fibroblasts. (H) Immunoblotting of the phosphoproteins (p-Akt/Akt and p-S6/S6) and LC3B (± Baf A1) in 143B cybrid cells bearing the m.8993T>G mutation showed no difference in protein phosphorylation and autophagic flux among cells with 0% (N0), 50% (N50) and 80% (N80) mutant loads (i, n = 3 independent experiments). PCR-RFLP was applied to measure the change in mutant load in 143B cybrid cells carrying the m.8993T>G mutation treated with LY or RP for 8 weeks, showing no significant effect on the mutant load of the m.8993T>G (ii, n = 3-4 independent biological samples). All data are represented as mean ± S.D. and were analysed by one/two-way ANOVA with Tukey’s multiple comparisons test (* *p* < 0.05, ** *p* < 0.01, *** *p* < 0.001, **** *p* < 0.0001).

**Tabel S1.**
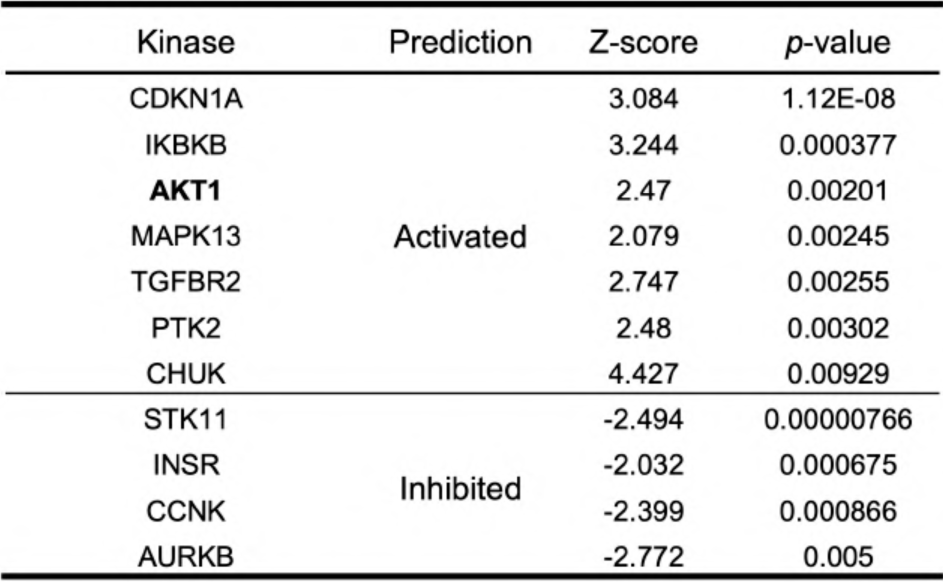
Upstream analysis for Kinase in patient fibroblasts by IPA.

**Table S2.** Upstream analysis for translation regulator in patient fibroblasts by IPA.

| Translation regulator | Prediction | Z-score | p-value |
| --- | --- | --- | --- |
| EIF4E | Inhibited | -2.169 | 0.000541 |
| EIF4G1 |  | -2.333 | 0.00245 |

**Table S3.**
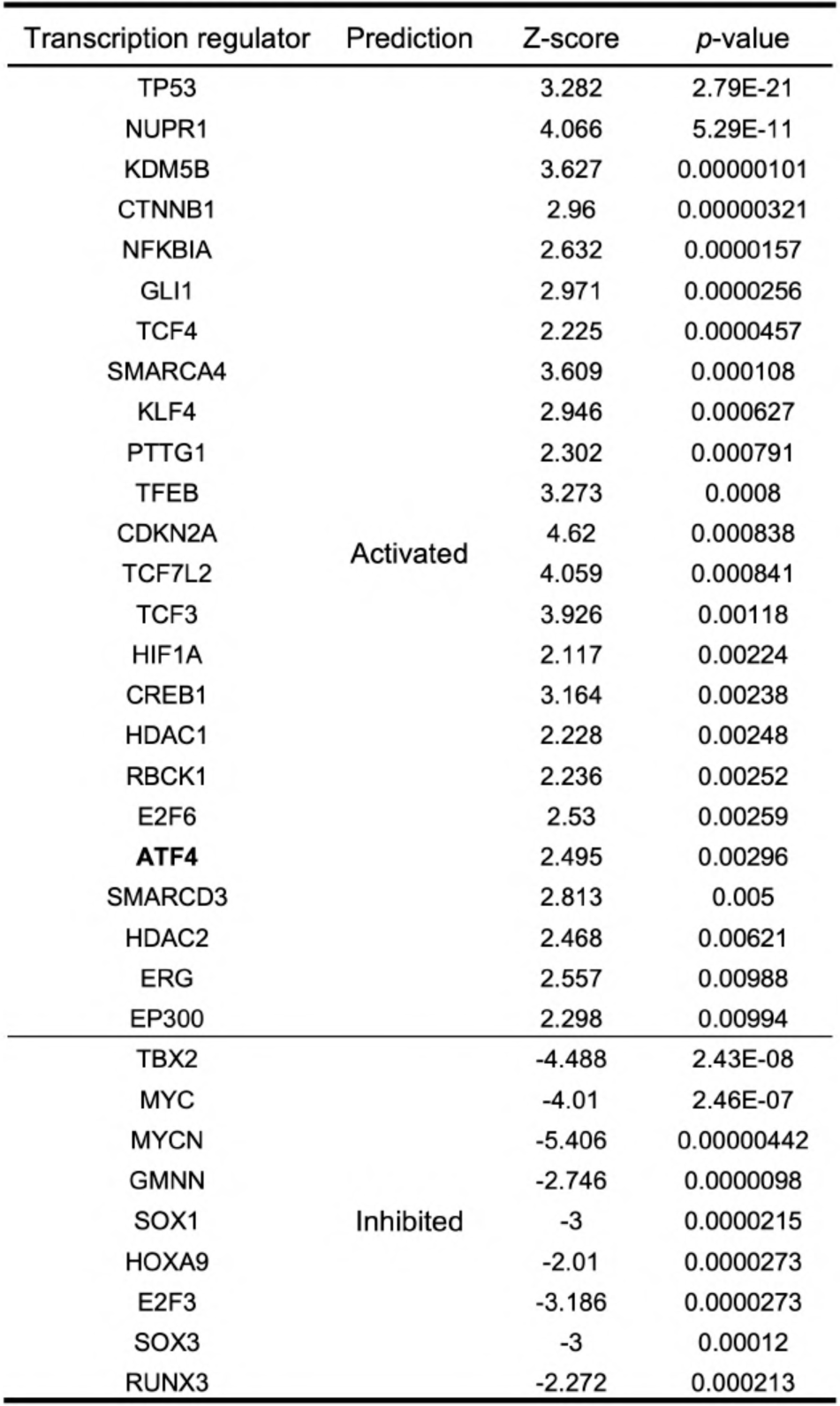
Upstream analysis for transcription regulators in patient fibroblasts by IPA.

| Transcription regulator | Prediction | Z-score | p-value |
| --- | --- | --- | --- |
| TP53 | Activated | 3.282 | 2.79E-21 |
| NUPR1 |  | 4.066 | 5.29E-11 |
| KDM5B |  | 3.627 | 0.00000101 |
| CTNNB1 |  | 2.96 | 0.00000321 |
| NFKBIA |  | 2.632 | 0.0000157 |
| GLI1 |  | 2.971 | 0.0000256 |
| TCF4 |  | 2.225 | 0.0000457 |
| SMARCA4 |  | 3.609 | 0.000108 |
| KLF4 |  | 2.946 | 0.000627 |
| PTTG1 |  | 2.302 | 0.000791 |
| TFEB |  | 3.273 | 0.0008 |
| CDKN2A |  | 4.62 | 0.000838 |
| TCF7L2 |  | 4.059 | 0.000841 |
| TCF3 |  | 3.926 | 0.00118 |
| HIF1A |  | 2.117 | 0.00224 |
| CREB1 |  | 3.164 | 0.00238 |
| HDAC1 |  | 2.228 | 0.00248 |
| RBCK1 |  | 2.236 | 0.00252 |
| E2F6 |  | 2.53 | 0.00259 |
| **ATF4** |  | 2.495 | 0.00296 |
| SMARCD3 |  | 2.813 | 0.005 |
| HDAC2 |  | 2.468 | 0.00621 |
| ERG |  | 2.557 | 0.00988 |
| EP300 |  | 2.298 | 0.00994 |
| TBX2 | Inhibited | -4.488 | 2.43E-08 |
| MYC |  | -4.01 | 2.46E-07 |
| MYCN |  | -5.406 | 0.00000442 |
| GMNN |  | -2.746 | 0.0000098 |
| SOX1 |  | -3 | 0.0000215 |
| HOXA9 |  | -2.01 | 0.0000273 |
| E2F3 |  | -3.186 | 0.0000273 |
| SOX3 |  | -3 | 0.00012 |
| RUNX3 |  | -2.272 | 0.000213 |

**Table S4.**
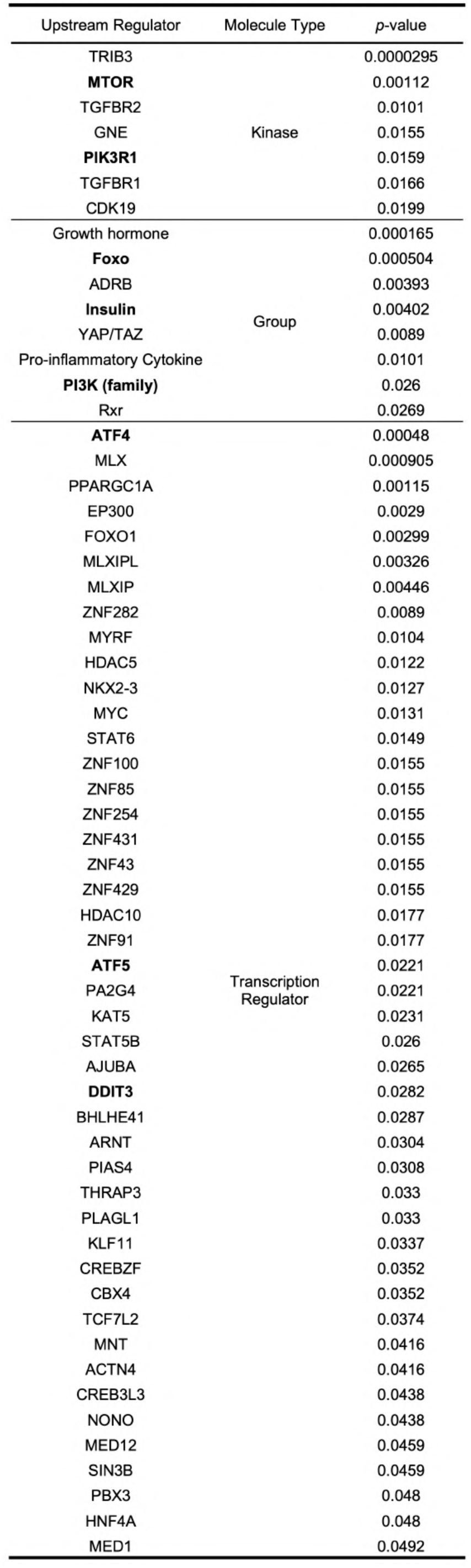
Upstream analysis using published RNA-seq dataset of patient biopsies by IPA.

## METHODS

### RESOURCE AVAILABILITY

#### Lead Contact

Further information and requests for resources and reagents should be directed to and will be fulfilled by the Lead Contact, Michael R Duchen.

#### Materials Availability

This study did not generate new unique reagents.

#### Data and Code Availability

Raw and processed RNA-seq data are accessible at the Gene Expression Omnibus under accession GSE175477. Other data including uncropped images of immunoblotting are available in the source and supplementary data.

### EXPERIMENTAL MODEL AND SUBJECT DETAILS

#### Cell Lines

The A549 cybrid cell line derived from the fusion of enucleated patient cells harbouring the m.3243A>G mtDNA mutation was a gift from Ian Holt (Biodondostia Research Institute, San Sebastián, Spain). The 143B cybrid cell lines with ∼50% and ∼80% of the m.8993T>G mtDNA mutant loads were gifts from the Minczuk lab (MRC Mitochondrial Biology Unit, Cambridge, UK). Control and patient fibroblasts bearing the m.3243A>G mtDNA mutation were obtained from the MRC Centre for Neuromuscular Disorders Biobank London. The patient fibroblasts with the m.3243A>G mutation were isolated from two female subjects (mother and daughter) at the age of 59 (patient 1) and 35 (patient 2) at the time biopsies were taken. For patient 1, the clinical symptoms included diabetes, myoclonus, sensorineural hearing loss, memory decline, myopathy, pigmentary retinopathy and bipolar affective disorder. For patient 2, the symptoms included diabetes, sensorineural deafness, cerebellar ataxia, myopathy, epilepsy, depression and cognitive impairment. These cell lines were cultured in Dulbecco’s modified Eagle’s medium (Gibco #10566016) supplemented with 10% fetal bovine serum (Gibco #16140071), and 1% Antibiotic-Antimycotic (Gibco #15240096) and incubated at 37 °C with 5% CO_2_.

#### Human muscle biopsies

The muscle biopsies were from a female patient with the m.3243A>G mutation (at the age of 55) and a matched healthy control. The study was approved by the Queen Square Research Ethics Committee, London (09/H0716/76). Informed consent was obtained from all participants.

### METHOD DETAILS

#### Cell culture and drug treatments for PI3K, Akt, or mTORC1 inhibition

A549 cybrid cells (WT and mutant) and 143B cybrid cells were passaged every 3-4 days at 80% confluence and trypsinised using 0.25% trypsin-EDTA (Gibco #25200056). Similarly, control and patient fibroblasts were passaged every week in the same way. Media with or without drugs were changed every 2-3 days. For drug treatments, rapamycin (Cayman Chemical #13346), LY294002 (Cayman Chemical #70920) GDC0941 (Cayman Chemical #11600) and MK2206 (Cayman Chemical #11593) were dissolved in DMSO (Sigma-Aldrich #D2650) at 10 mM stock concentrations. For assays to assess autophagy, chloroquine (Cayman Chemical #14194) was dissolved in PBS at 50 mM for stock, while Bafilomycin A1 (Cayman Chemical #11038) was dissolved in DMSO for stock at 200 μM. Drugs were subsequently diluted to their working concentrations in media. Heteroplasmy level in the A549 cybrid cells, the 143B cybrid cells and the fibroblasts of patient 1 without treatments did not change significantly throughout the study as assessed by ARMS-qPCR or PCR-RFLP.

#### Measurements for mtDNA mutations

Levels of mtDNA mutation were detected using PCR-restriction fragment length polymorphism (RFLP)^16, 48^ or allele refractory mutation system (ARMS)-based quantitative PCR (qPCR) analysis^17^. DNA extractions from frozen cells were performed using the DNeasy Blood & Tissue Kit (Qiagen #69506). Concentrations of DNA samples were quantified using NanoDrop. All primer pairs used can be found in Table S5. For PCR-RFLP, samples (20 ng/µl, 2 μl) were mixed with the master mixes containing 1 μl primers (10 μM each) and reagents of GoTaq G2 (Promega #M7845) and the final volume was 25 μl. After 30 thermocycles, amplified PCR products (8 μl) were further digested by ApaI (Promega #R6361) for the m.3243A>G and HapII (Promega #R6311) for the m.8993T>G and analysed by a 2% agarose gel with ethidium bromide. Densitometric analysis was performed by Fiji^49^. For ARMS-qPCR, samples were diluted to 0.4 ng/µl. ARMS primer working solutions (5 μM, 1 μl each) and SYBR Green JumpStart Taq ReadyMix (Sigma-Aldrich #S4438) were added together as master mixes for mutant and wild-type genes. DNA samples (3 μl) and master mixes (7 μl) were pipetted into a 96-well PCR plate (Bio-Rad #MLL9651) and PCR amplification performed using the CFX96 Touch Real-Time PCR Detection System (Bio-Rad). Each sample has three technical replicates. Mutant heteroplasmy level (%) was calculated using 1/[1 + (1/2) ^ΔCT^] × 100%, where ΔC_T_ (cycle threshold) = CT_wild-type_ – CT_mutant_.

ARMS-qPCR is a quantitative way to measure mutant loads. In contrast, PCR-RFLP is relatively a qualitative way to detect mtDNA mutations. Due to an intrinsic problem of PCR-RFLP, part of the PCR products are hybrids which have a strand of wild-type (WT) and a strand of mutant, because a single nucleotide mutation is not strong enough to avoid their annealing. Moreover, restriction enzymes cannot recognise and cleavage these hybrids of WT/mutant double strand DNA, which means that in the “WT” band, part of the PCR products are hybrids. Thus, the PCR-RFLP largely underestimates the mtDNA mutant loads, which is the reason why there are weaker hands of the mutant than which of the WT.

#### Quantification of relative mtDNA copy number

The relative mtDNA copy number of cells was determined using quantitative PCR with primers for the mtDNA tRNA^Leu(UUR)^ and with primers for the nuclear B2-microglobulin as previously described^50^. Standard 96 well PCR plate with optically clear sealing film and CFX96 Real-Time PCR Detection System (Bio-Rad) were used. PCR mix consists of 2 μl of template DNA (3 ng/μl), 2 μl of primer pair (final concentration of 400 nM), 12.5 μl of SYBR Green JumpStart Taq ReadyMix (Sigma-Aldrich) and 8.5 μl of DNase/RNase free H_2_O. The thermal cycling conditions were as follow: 50 °C for 2 min, 95 °C for 10 min and then 40 cycles of 95 °C for 15 s and 62 °C for 1 min. Each sample has three technical replicates. The following equation was used to determine the relative mitochondrial DNA content, 2 x 2^ΔCT^, where ΔC_T_ is nuclear DNA CT value subtracted by mtDNA CT value.

#### The mitochondrial oxygen consumption rate

Measurements of aerobic respiration and glycolysis were conducted with the Seahorse Bioscience XFe-96 bioanalyzer using the Seahorse XF Cell Mito Stress Test Kit (Agilent #103015-100). Cells were seeded on XF96 cell culture microplates (Agilent #102416-100) two days before the experiment (A549 cybrid cells, 1×10^4^ cells/well; Fibroblasts, 2×10^4^ cells/well). On the day of the experiment, the culture medium was replaced with Seahorse XF Base medium (Agilent #103334-100) supplemented with 1 mM pyruvate (Gibco #11360070), 2 mM glutamine (Gibco #25030081) and 10 mM glucose (Gibco #A2494001) and incubated for 30 min at 37 °C in a CO_2_-free incubator before loading into the Seahorse Analyzer. After measuring basal respiration, the drugs oligomycin (5 µM), FCCP (1 µM, 2 µM), and rotenone/antimycin A (0.5 µM/0.5 µM) were added to each well in sequential order. Data were analysed using the XF Cell Mito Stress Test Report Generator. After the assay, cells were stained with Hoechst 33342 (5 µM; Thermo Scientific #62249) for 30 min. ImageXpress was then used to count the numbers of cell nuclei (cell numbers) in each well. The normalisation of the experiments is based on the relative cell numbers obtained.

#### Mitochondrial membrane potential (Δψ_m_)

Cells were seeded (3 x 10^4^/well for A549 cybrid cells; 1 x 10^4^/well for fibroblasts) in glass-bottom 24-well plates, 2-3 days before imaging. Cells were washed twice with the recording medium, which was phenol red-free DMEM (Gibco #A1443001) with 10 mM glucose, 1 mM glutamine, 10 mM HEPES, adjusted to pH 7.4; and then incubated with 25 nM tetramethylrhodamine methyl ester (TMRM) for 30 min at 37 °C. Cells were imaged with an LSM 880 (Carl Zeiss) confocal microscope using Fluar 63x/1.40 oil immersion objective lens at 37 °C. TMRM was excited with a 561 nm Argon laser with an output power of 0.2 mW. MBS 488/561 was used as a beam splitter and emitted fluorescence collected at 564-740 nm. Images were acquired using Zen Black software (Carl Zeiss) and fluorescence intensity was quantified using Fiji with the same threshold across all samples.

#### ROS measurements

Rate of general ROS production was assessed using dihydroethidium (DHE, Invitrogen #D11347), an intracellular superoxide indicator which is oxidised by superoxide to ethidium which fluoresces red. As for mitochondrial ROS production, MitoSOX (Invitrogen #M36008) was used. One day before the experiment, cells were seeded in a glass-bottom 96-well plate (SensoPlate #655892) at a density of 2 x 10^4^ cells per well. On the day of the experiment, cells were washed twice with PBS and incubated with 5 µM DHE or MitoSOX in recording medium (phenol red-free DMEM, 10 mM glucose, 1 mM glutamine, 10 mM HEPES, adjusted to pH 7.4). Measurements of fluorescence intensity were taken at intervals of 5 min at 37 °C using the CLARIOstar microplate reader (excitation/emission = 518/606 nm for DHE excitation/emission = 510/580 nm for MitoSOX). The total incubation time for DHE is 1 h and 30 min for MitoSOX. The slope of fluorescence intensity was calculated (Fig. S1C) and represents the rate of ROS production.

#### Glucose uptake using 2-NBDG

A fluorescent deoxyglucose analogue, 2-NBDG (Invitrogen #N13195), was employed as a probe to measure rates of glucose uptake by cultured cells. Cells were seeded in a glass-bottom 96-well plate as ROS measurements. On the day of experiments, cells were firstly washed with glucose-free recording media twice and incubated with the glucose-free media for 1 h. After media aspiration, recording media with 2-NBDG (100 µg/ml) were then added into wells and incubated for 30 min. After the incubation, cells were then again washed with glucose-free recording media twice and fluorescence intensity were measured using the CLARIOstar microplate reader (excitation/emission = 467/542 nm).

#### Medium pH values

Medium pH values were measured based on the ratiometric property of phenol red, a common pH indicator in media. The absorbance of phenol red change in response to changing pH. Cells were seeded in 96-well plates at a density of 1 x 10^4^ cells per well with 200 µl media and cultured for 2 days. On the day of experiments, media of each well were then transferred to a new 96-well plate and measured the absorbance of phenol red at 443 and 570 nm immediately. The higher the absorbance ratios of 443 to 570 nm, the more acidic the media.

#### Glucose and lactate concentrations in media using CuBiAn

Media for pH measurements were then directly used for quantifying glucose and lactate concentrations in media. Samples and fresh media were measured using the CuBiAn HT-270 biochemistry analyser (Optocell technology) with its Glucose (#200106) and Lactate Assay Kits (#200115) according to manufacturer instructions.

#### Blue native gel electrophoresis (BNGE) and In Gel activity assays

Mitochondria were isolated from cultured fibroblasts and cell lines according to the method described earlier (ref). Digitonin-solubilized mitochondria proteins (100 µg) were separated on pre-cast 3%-12% gradient blue native gels (Invitrogen #BN1001) according to manufacturer’s instructions. After electrophoresis, the gels were electroblotted onto PVDF membrane (Millipore #IPVH00010) and probed with anti-OxPhos antibody cocktail (1:1000, Invitrogen #45-8199), anti-SDHA (1:1000, Abcam #ab137040) and anti-ATP5A (1:1000, Abcam #ab14748). The enzymatic activity of different OxPhos complexes was determined by in-gel assays. For CIV+CI activity, the gels were incubated first in CIV substrate (50 mg diaminobenzidine and 100 mg cytochrome c in 50 mM phosphate buffer, pH 7.4) until brown signal was observed and then incubated in CI substrate (0.1 mg/ml NADH, 2.5 mg/ml Nitrotetrazolium Blue chloride in 100 mM Tris–HCl, pH 7.4) until blue signal appeared. The reaction was stopped with 10% acetic acid and the gels were washed and scanned.

#### SDS-PAGE and immunoblotting

For immunoblotting, cells were seeded 1-day prior experiments (60 mm plates for the A549 cybrid cells and 10 cm plates for fibroblasts). To assess autophagic flux, cells were replenished with regular media for 1 h to prevent starvation-induced autophagy. After 1 h, treatment conditions resumed either with or without 50 μM chloroquine for 6 h. For other experiments, cells were then washed with ice-cold PBS once and lysed using 150-300 µl RIPA buffer (Sigma-Aldrich #R0278) with one cOmplete™ Protease Inhibitor Cocktail (Roche #4693116001) tablet and one PhosSTOP Phosphatase Inhibitor Cocktail (Roche #4906837001) tablet. Cells were then scraped and centrifuged at 16,000 g at 4 °C for 30 min. Protein concentration in the supernatant was quantified using the Pierce BCA Assay Kit (Thermo Scientific #23227). For immunoblotting, 30 µg of protein samples in NuPAGE 4x LDS Sample Buffer (Invitrogen #NP0007) and 2% β-mercaptoethanol (Sigma-Aldrich #63689) were boiled at 99°C for 5 min. Proteins were separated on 4-12% NuPAGE Bis-Tris polyacrylamide gels (Invitrogen #NP0335) or 12% Bis-Tris gels (#NW00122) immersed in MOPS running buffer (Invitrogen #NP0001) and transferred onto PVDF membranes (Millipore #IPFL00010). Membranes were then incubated in Intercept (TBS) Blocking Buffer (Li-COR Biosciences #927-60001) for 1 h at room temperature. After addition of primary antibodies diluted in the blocking buffer with 0.1% Tween-20, membranes were incubated overnight at 4°C on a shaker. Subsequently, membranes were incubated with appropriate secondary antibodies (Li-COR Biosciences; 1:10000; IRDye® 680RD Goat anti-Mouse IgG, #926-68070; IRDye® 800CW Goat anti-Rabbit IgG, #926-32211) for 1 h at room temperature before signals were developed with the LiCOR Odyssey CLx system. Details of all the antibodies used in this study can be found in the Key Resources Table. Following is the antibodies and dilutions for immunoblotting: Anti-PC (1:1000, Novus Biologicals #NBP1-49536), anti-phospho-PDHA (1:1000, Millipore #AP1062), anti-PDHA (1:1000, Invitrogen #45-6600), anti-p-Akt (1:1000, Cell Signaling Technology #9271), anti-Akt (1:3000, Cell Signaling Technology #9272), anti-p-S6 (1:3000, Cell Signaling Technology #4858), anti-S6 (1:3000, Cell Signaling Technology #2217), anti-p-AMPK (1:1000, Cell Signaling Technology #2532), anti-AMPK (1:3000, Cell Signaling Technology #2532), anti-LC3B (1:3000, Cell Signaling Technology #3868), and anti-β-actin (1:10000, Santa Cruz Biotechnology #sc-47778).

#### Live-cell imaging for cell growth and death using the Incucyte platform

Cells were seeded at a density of 2,000 cells/well in a 96-well plate one-day prior imaging with regular cell culture media. For cell growth/death in a variety of nutrient conditions, cells were washed with PBS twice and then DMEM (Gibco #11966025 or #11960044) with different concentration of glucose, galactose (Sigma-Aldrich #G5388) or glutamine and 10% dialysed FBS (Sigma-Aldrich #F0392) were added. For drug treatments, media were then replaced with fresh media with/without drugs along with 20 nM SYTOGreen Nucleic Acid Stain (Invitrogen #S7572) just before imaging. Images were acquired using the Incucyte with 20X objective every 2 h for a 3-4 day period. Cell confluency and cell death numbers were analysis on Incucyte software and sequentially extracted. Rates of cell growth were analysed by fitting the growth curve to the exponential cell growth model (Fig. S1E) using the solver in Excel, while cell death was obtained by simply normalising the numbers of dead cells (SYTOGreen stained) to total cell confluency.

#### Measurements of cytosolic NADH:NAD^+^ using SoNar

The genetically encoded probes, SoNar for NADH:NAD^+^ ratio, were developed by and obtained from the Yang’s Lab (Chinese Academy of Sciences)^18^. For A549 cybrid cells, the cells were seeded at the density of 4 x 10^4^/well in glass-bottom 24-well plates two days before transfection. Lipofectamine 3000 (Invitrogen #L3000001) was used for transfection according to the manufacturer’s protocol. Specifically, we used 1 µg DNA and the ratio of p3000 to the 3000 reagent was 2:1. The cells were imaged two days after transfection in recording media with the LSM 880 microscope using 20x/0.8 objective lens at 37°C. Both probes were excited at 405 and 488 separately and emitted fluorescence longer than 535 nm was collected. For fibroblasts, Human Dermal Fibroblasts Nucleofector Kit (Lonza #VPD-1001) was used for transfection according to the manufacturer’s protocol. Specifically, we used 2.5 µg DNA for 5 x 10^5^ cells. Similarly, fibroblasts were imaged two days after transfection with the same conditions as A549 cybrid cells. Images were acquired using Zen Black software (Carl Zeiss) and analysed using Fiji. Regions of interest (ROI) were generated by thresholding and binarizing images and average fluorescence intensity in the ROIs for each channel were then measured. Ratios between the signal excited at 405 nm and 488 nm were calculated.

#### Measurement of autophagy and mitophagy using mCherry-EGFP-LC3 and mt-Keima/COX8-EGFP-mCherry reporters, respectively

The autophagy reporter, mCherry-EGFP-LC3 (Addgene plasmid #22418), and mitophagy reporter, mt-Keima (Addgene plasmid #56018) and COX8-EGFP-mCherry (Addgene plasmid #78520), were purchased from Addgene. The procedure of transfection for A549 cybrid cells and patient fibroblasts was similar to SoNar. Imaging was performed with the LSM 880 microscope using 63x/1.40 oil immersion objective lens at 37 °C. The mCherry-EGFP-LC3 was excited at 488 and 561 nm separately and emitted fluorescence 500-580 and longer than 600 nm was collected. Images were acquired using Zen Black software (Carl Zeiss). Numbers and area of fluorescent particles for each channel were quantified using Fiji by thresholding images. For mt-Keima, following transfection, the cells were treated as indicated for 24h and imaged via two sequential excitations (458 nm, green; 561 nm, red) using a 570-to 695-nm emission range. The laser power was set at the minimum output to allow the clear visualization of the mt-Keima signal and was separately adjusted for each experimental condition. At least 10 z-stacks with 0.45 um thickness were acquired per sample in an experimental set. LysoTracker Green DND-26 (Invitrogen #L7526) was co-imaged using a 488 nm excitation and a 495-550 nm emission filter, where indicated. The ratio (high F_543_/F_458_ ratio area/total mitochondrial area) was used as an index of mitophagy. Ratio (F_543_/F_458_) images were generated using the Ratio Plus plugin in Fiji. High (F_543_/F_458_) ratio areas and total mitochondrial area were binarized, segmented and quantified in Fiji. Quantitative analysis of mitophagy in COX8-EGFP-mCherry-transfected A549 cybrid cells was also assessed using a flow cytometry-based approach in which an arbitrary threshold of red fluorescence signal was used to indicate a mitophagic positive cell. Following transfection, cells were treated with either DMSO or different drugs as indicated. After 24 h. of treatment, cells were trypsinized, washed once with DPBS and then resuspended into 1 ml of DPBS prior to analysis using a BD Accuri C6 flow cytometer. Cells were excited with a 488 nm laser with emission assessed simultaneously using a 533/30 nm (FL1 detector) and a 670 LP filter (FL3 detector). Cells with high FL3/FL1 ratio were selected and their proportion was calculated.

#### Metabolomics

Control and patient fibroblasts were plated 24 hrs before the experiment in 10 cm dishes (1 x 10^6^ cells/dish). On the day of experiments, the medium was replaced with phenol red-free DMEM with 1 mM glutamine, 10% dialysed FBS (prepared as described in Yuneva et al., 2007) and 10 mM [U-^13^C]-glucose (Goss Scientific #CLM-1396-1). After 24 h incubation, cells were washed once with ice-cold PBS. Metabolites were extracted with adding 500 µl of ice-cold methanol, scraping, and transferring into tubes on ice. Dishes were then subsequently washed with 250 µl methanol and 250 µl water containing 2.5 nM of each nor-leucine and scyllo-inositol per sample as internal standards. Finally, 250 µl of chloroform was added and samples were sonicated for 3 rounds (8 min each) at 4 °C. After centrifugation (18,000 g for 10 min at 4 °C), the upper polar fraction was collected and used for GC-MS analysis, while he protein pellets were dried, lysed using 62.5 mM Tris buffer (pH 6.8) containing 2% SDS and used for protein quantification using BCA assay (Abcam). Briefly, the polar fraction was washed twice with methanol, derivatized by methoximation (Sigma, 20 μl, 20 mg/ml in pyridine) and trimethylsilylation (20 μl of N,O-bis(trimethylsilyl) trifluoroacetamide reagent (BSTFA) containing 1% trimethylchlorosilane (TMCS), Supelco), and analysed on an Agilent 7890A-5975C GC–MS system. Splitless injection (injection temperature 270 °C) onto a 30 m + 10 m × 0.25 mm DB-5MS + DG column (Agilent J&W) was used, using helium as the carrier gas, in electron ionization (EI) mode. The initial oven temperature was 70 °C (2 min), followed by temperature gradients to 295 °C at 12.5 °C/min and then to 320 °C at 25 °C/min (held for 3 min). Data analysis and peak quantifications were performed using MassHunter Quantitative Analysis software (B.06.00 SP01, Agilent Technologies). The level of labelling of individual metabolites was corrected for the natural abundance of isotopes in both the metabolite and the derivatization reagent. Abundance was calculated by comparison to responses of known amounts of authentic standards^52^. The data were further processed and visualised by MetaboAnalyst 5.0, using the Pathway Analysis module (data were normalised to controls and scaled by the standard deviation of each variable) with the library of human SMPDB^53^.

#### RNA-sequencing

Fibroblasts for RNA-sequencing were plated in 10 cm dishes with regular media (1 x 10^6^ cells/dish) one day before RNA extraction. RNA extractions were performed using the RNeasy Mini Kit (Qiagen #74104). Concentrations and quality of RNA samples were quantified using NanoDrop (total RNA >250 ng, A_260_/A_280_ = 1.8-2.1, A_260_/A_230_ > 1.7). RNA-sequencing was then performed at UCL Genomics and analysed by the SARTools R package^54^. The resulting datasets were further analysed by Ingenuity Pathway Analysis (IPA, Qiagen) with a threshold of FDR < 0.05 and visualised by Morpheus (https://software.broadinstitute.org/morpheus) for heatmaps and NetworkAnalyst 3.0^55^ for volcano plots and principal component analysis.

#### Immunohistochemistry

For immunohistochemistry, muscle biopsies were fixed in phosphate buffered saline (PBS) and fixed in 4% paraformaldehyde (PFA), made in PBS, for 4-8 h at room temperature (RT). Following fixation, tissues were cryoprotected in 30% sucrose in Diethyl Pyrocarbonate (DEPC) treated PBS, embedded and frozen in a mixture of 15% sucrose /50% Tissue-Tek OCT (Sakura Finetek), and sectioned in the coronal plane at 20 µm using a Cryostat (Bright Instruments). Patient muscle biopsies sections were washed in PBS, blocked in a solution of 5% normal goat serum (Merck KGaA) (v/v) containing 0.1% Triton X-100 (v/v) (Merck KGaA) in PBS at RT for 2 h. They were first incubated in primary antibodies at RT overnight. The following antibodies were used: pan-AKT (1:200, Cell Signaling Technology #2920), phospho-AKT (p-AKT; 1:200, Abcam #ab81283), S6 (1:200), and phospho-S6 (1:200). Following incubation in primary antibodies, sections were washed in PBS, incubated in biotinylated anti-species secondary antibodies (1:250; Vector Laboratories) for 2 h. Sections were washed and incubated with bisbenzimide (10 min in 2.5 μg/ml solution in PBS; Merck KGaA). Images were collected using an SP2 Leica confocal microscope (Leica Microsystems, UK). Sequential images were subsequently reconstructed using Metamorph imaging software (Universal Imaging Corporation, West Chester, PA).

#### Immunofluorescence

Fibroblasts were seeded at a density of 5 x 10^4^ cells/well in 24 well plates on 10 mm coverslips. After 24h incubation, cells were treated with LY or RP for 4 weeks and 8 weeks respectively. After the long-term treatment, the cells were washed thrice with 1X PBS and fixed in 4% paraformaldehyde for 15 mins at room temperature and permeabilised with 0.1% Triton-X 100 for 15 min in PBS. The cells were then washed and incubated with anti-MTCOI antibody (1:100, Abcam #ab14705) in 3% BSA for 1h at the room temperature followed by incubation with Alexa Fluor 488-conjugated secondary antibody for 1h at the room temperature. After antibody labelling, the coverslips were mounted on a glass slide using ProLong™ Gold Antifade mountant with DAPI and imaged using the confocal microscope as described above.

#### TaqMan SNP genotyping for Single cell qPCR

Primers for distinguishing WT and the m.3243A>G mutant mtDNA were designed by submitting the region of the m.3243A>G mutation (∼120 bp) to the website of online Custom TaqMan® Assay design tool (https://www.thermofisher.com/order/custom-genomic-products/tools/genotyping/) The Custom TaqMan SNP Genotyping Kit (Applied Biosystems #4332073) was ordered and prepared according to manufacturer instructions. For single cell sorting, cells were resuspended (1 x 10^6^ cells/ml for A549 cybrid cells; 5 x 10^5^ cells/ml for fibroblasts) in 1 ml PBS with 1% FBS, filtered through 70 μm mesh and then kept on ice. Sorting cells to 96-well PCR plates using BD FACS Aria Fusion was performed by Flow Cytometry Core Facility, Division of Medicine at UCL. Cells in 96-well plates were lysed with 2 μl of 0.2% Triton X-100 and kept in −80 °C immediately. For qPCR, TaqPath ProAmp Master Mix (Applied Biosystems #A30866) was mixed with primers of TaqMan SNP genotyping and added into 96-well PCR plates to reach a final volume of 10 μl in each well. Thermo cycles were set according to manufacturer instructions. Mutant loads were determined by the ratio of mutant to total fluorescent intensity.

#### Quantification and statistical analysis

All statistical analyses, unless otherwise stated in figure legends, were carried out using GraphPad Prism 8. To compare means between two groups, a two-tailed unpaired t-test was used for normally distributed data. One/two-way ANOVA with Tukey’s multiple comparisons test was performed for multi-group (at least three) comparisons. Data are presented as graphs displaying mean ± S.D., of at least three independent biological replicates. Means of control samples on immunoblotting or immunofluorescence are typically centred at one (or 100%) to ensure easier comparisons unless otherwise noted. Differences were only considered to be statistically significant when the *p* value was less than 0.05. Estimated *p* values are either stated as the actual values or denoted by * *p* < 0.05, ** *p* < 0.01, *** *p* < 0.001, **** *p* < 0.0001.

For detailed processing and statistical analysis of RNA sequencing and metabolomics, please find them in the previous section with the same subheadings.

No statistical method was used to predetermine sample size, and replicates are shown in Figure legends. The investigators were not blinded to allocation during experiments and outcome assessment.

## ACKNOWLEDGEMENTS

We thank Alice Giustacchini (UCL ICH) for help with single cell analyses, Stephanie Carrington (Division of Neuropathology, UCL IoN) for sectioning the muscle biopsies, Ash Merve (Consultant Neuropathologist, Division of Neuropathology, UCL IoN and Department of Histopathology, Camelia Botnar Laboratory, Great Ormond Street Hospital) for reviewing the biopsy and taking H&E staining images, and Metabolomics STP of the Francis Crick Institute for assistance with GC-MS analysis. We also acknowledge The MRC Centre for Neuromuscular Diseases Biobank (supported by the National Institute for Health Research Biomedical Research Centres at Great Ormond Street Hospital for Children NHS Foundation Trust and at University College London Hospitals NHS Foundation Trust and University College London) for providing all patient-derived cells used in this study. RDSP is supported by a Medical Research Council (UK) Clinician Scientist Fellowship (MR/S002065/1) and a Medical Research Council (UK) strategic award to establish an International Centre for Genomic Medicine in Neuromuscular Diseases (ICGNMD) (MR/S005021/1). GS is supported by Telethon GGP16026 and CRUK Pioneer Award C28472/A29264. MY is supported by the Francis Crick Institute which receives its core funding from Cancer Research UK (FC001223), the UK Medical Research Council (FC001223), and the Wellcome Trust (FC001223). GEV is supported by the National Agency for Research and Development (ANID) / Scholarship Program / DOCTORADO BECAS CHILE/2019 – 7220052 We also thank Action Medical Research for funding the early stages of this work and the Ministry of Education, Taiwan for funding CYC’s PhD by the Government Scholarship to Study Abroad.

